# A positional and combinatorial regulatory code for alternative splicing

**DOI:** 10.64898/2026.06.10.729476

**Authors:** Mina Yoshida, Masahiko Ajiro, Hiroki Ueda, Kazuki Nishimura, Ryoichi Maenosono, Mayumi Hanzawa, Natsuko Shinohara, Marimu Sakumoto, Syuzo Kaneko, Ryuji Hamamoto, Rinshi S. Kasai, Genta Nagae, Hirotaka Matsui, Atsushi Iwama, Hiroyuki Aburatani, Shungo Adachi, Asuka Kawachi, Akihide Yoshimi

**Affiliations:** Division of Cancer RNA Research, National Cancer Center Research Institute, Tokyo, Japan; Health and Disease Omics Center, Chiba University, Chiba, Japan; Division of Medical AI Research and Development, National Cancer Center Research Institute, Tokyo, Japan; Division of Advanced Bioimaging, National Cancer Center Research Institute, Tokyo, Japan; Genome Science & Medicine Laboratory, Research Center for Advanced Science and Technology, The University of Tokyo, Tokyo, Japan; Department of Laboratory Medicine, National Cancer Center Hospital, Tokyo, Japan; Division of Stem Cell & Molecular Medicine, The Institute of Medical Science, The University of Tokyo, Tokyo, Japan; Division of Proteomics, National Cancer Center Research Institute, Tokyo, Japan

**Author notes:** Correspondence: Masahiko Ajiro, PhD, Senior Research Scientist, Division of Cancer RNA Research National Cancer Center Research Institute, 5-1-1 Tsukiji, Chuo-ku, Tokyo, 1040045, Japan, Akihide Yoshimi, MD, PhD, Chief, Division of Cancer RNA Research National Cancer Center Research Institute, 5-1-1 Tsukiji, Chuo-ku, Tokyo, 1040045, Japan.

## Abstract

Alternative pre-mRNA splicing generates extensive transcript diversity, yet the regulatory code that determines how splicing decisions are encoded across the transcriptome remains poorly defined. Splicing outcomes are controlled by combinatorial RNA-binding protein (RBP) interactions and positional context, but how these features are integrated at the transcriptome scale remains unclear. CLIP-based approaches have mapped RBP binding, but directly comparable endogenous maps across multiple RBPs are lacking, limiting inference of global regulatory principles. Here we introduce SCALE-CLIP, an endogenous CLIP framework that integrates CRISPR-Cas9-mediated epitope tagging with long-read-guided read attribution to generate directly comparable RBP binding maps across splicing-regulatory factors. Applied to 23 RBPs, SCALE-CLIP expanded endogenous RBP coverage and, across benchmarked shared factors, increased peak recovery by a median of 12.2-fold relative to ENCODE eCLIP while preserving specificity and reproducibility. We define a transcriptome-wide positional and combinatorial code for alternative splicing, in which binding position is a primary determinant of regulatory outcome: SRSF binding within alternative exons promotes inclusion, whereas binding on flanking exons drives exon skipping. Higher-order SRSF occupancy further tunes this code, buffering exon inclusion when centered on alternative exons but reinforcing repression when distributed across flanking exons. We also show that m^6^A provides an epitranscriptomic layer that locally enhances SRSF binding and is associated with increased exon inclusion. Together, these results establish a multi-layered RNA-binding logic in which binding position, combinatorial RBP architecture and RNA modification jointly shape splicing outcomes, providing a framework for rational interpretation and modulation of alternative splicing.

## Introduction

Alternative pre-mRNA splicing is a fundamental source of proteomic diversity in eukaryotes^1^. By selectively including or excluding exons, splicing generates multiple mRNA isoforms from a single gene, thereby shaping gene expression programs across development, physiology and disease^2,3^. Although splicing decisions are governed by complex regulatory inputs, the underlying regulatory code that determines how these decisions are encoded across the transcriptome remains poorly defined.

At the molecular level, splicing outcomes are determined by integration of multiple regulatory layers, including combinatorial RNA-binding protein (RBP) networks, binding position, co-transcriptional RNA–protein interactions, and local RNA sequence and structural context^4–6^. This regulatory logic is inherently difficult to resolve because splicing outcomes often reflect coordinated or competing contributions from multiple factors acting on the same transcript. An additional layer of complexity is provided by epitranscriptomic RNA modifications, which can influence RNA processing by altering RNA structure, modulating RBP accessibility, or directly affecting splice-site recognition^7–9^. However, how these regulatory layers are quantitatively integrated into a unified framework across transcripts under endogenous conditions remains unclear. In particular, directly comparable binding maps across major splicing-regulatory RBPs are still limited, preventing causal inference and limiting modelling of how binding organization, positional context, and modification state together encode splicing outcomes.

Transcriptome-wide mapping of RBP binding by crosslinking and immunoprecipitation (CLIP) and related approaches has provided key insights into splicing regulation^10–15^. However, these approaches remain limited in several important respects. They depend on antibody performance, restricting factor coverage and complicating quantitative comparisons across RBPs, and overexpression-based strategies can perturb native expression and interaction networks, confounding inference under physiological conditions^16–20^. In addition, conventional short-read CLIP data are intrinsically difficult to interpret in repetitive or low-uniqueness regions, reducing recoverable signal in repeat-rich regulatory landscapes^14,21,22^. Together, these limitations preclude systematic and directly comparable analysis of RBP binding organization across factors, preventing reconstruction of a unified regulatory code linking RBP binding architecture to splicing outcomes at the transcriptome scale.

To address these limitations, we developed SCALE-CLIP (Standardized, Comparable, Attribution-guided, Long-read-integrated enhanced CLIP-sequence), an endogenous eCLIP framework that combines CRISPR–Cas9-mediated epitope tagging with long-read-guided read attribution (LRA) to improve CLIP-read assignment in low-uniqueness regions and enable systematic comparison of RBP-binding landscapes across splicing-regulatory factors.

Using SCALE-CLIP, we systematically mapped the binding landscapes of splicing-regulatory RBPs and uncovered a transcriptome-wide positional and combinatorial code for alternative splicing. This analysis revealed a positional rule for SRSF-dependent splicing, in which binding within alternative exons and flanking exons has opposite regulatory effects, and showed that this rule is further tuned by SRSF co-occupancy and m^6^A modification. Together, these findings establish a multi-layered view of splicing regulation in which RBP position, higher-order occupancy and RNA modification jointly shape exon choice.

## Results

### SCALE-CLIP improves sensitivity and comparability of RBP mapping

To systematically detect RBP binding under endogenous conditions, we developed SCALE-CLIP, a CLIP framework that couples CRISPR-Cas9-mediated epitope tagging with LRA (Fig. 1a, Extended Data Fig. 1a–c, and Methods). We applied SCALE-CLIP to 23 splicing-regulatory RBPs, including seven factors with available ENCODE eCLIP datasets and 16 additional factors lacking high-quality ENCODE profiles in K562 cells. Tagged proteins showed expected expression, predominantly nuclear localization^23,24^, and low background RNA recovery, supporting the specificity and integrity of the SCALE-CLIP workflow (Extended Data Fig. 1d–g). We then implemented an LRA strategy, which uses long CLIP reads to select short-read alignments supported by transcript structure and splice-junction evidence, thereby expanding the recoverable binding landscape (Fig. 1b, Extended Data Fig. 2a-e, see Methods for details)^14,25^.

**Fig. 1.**
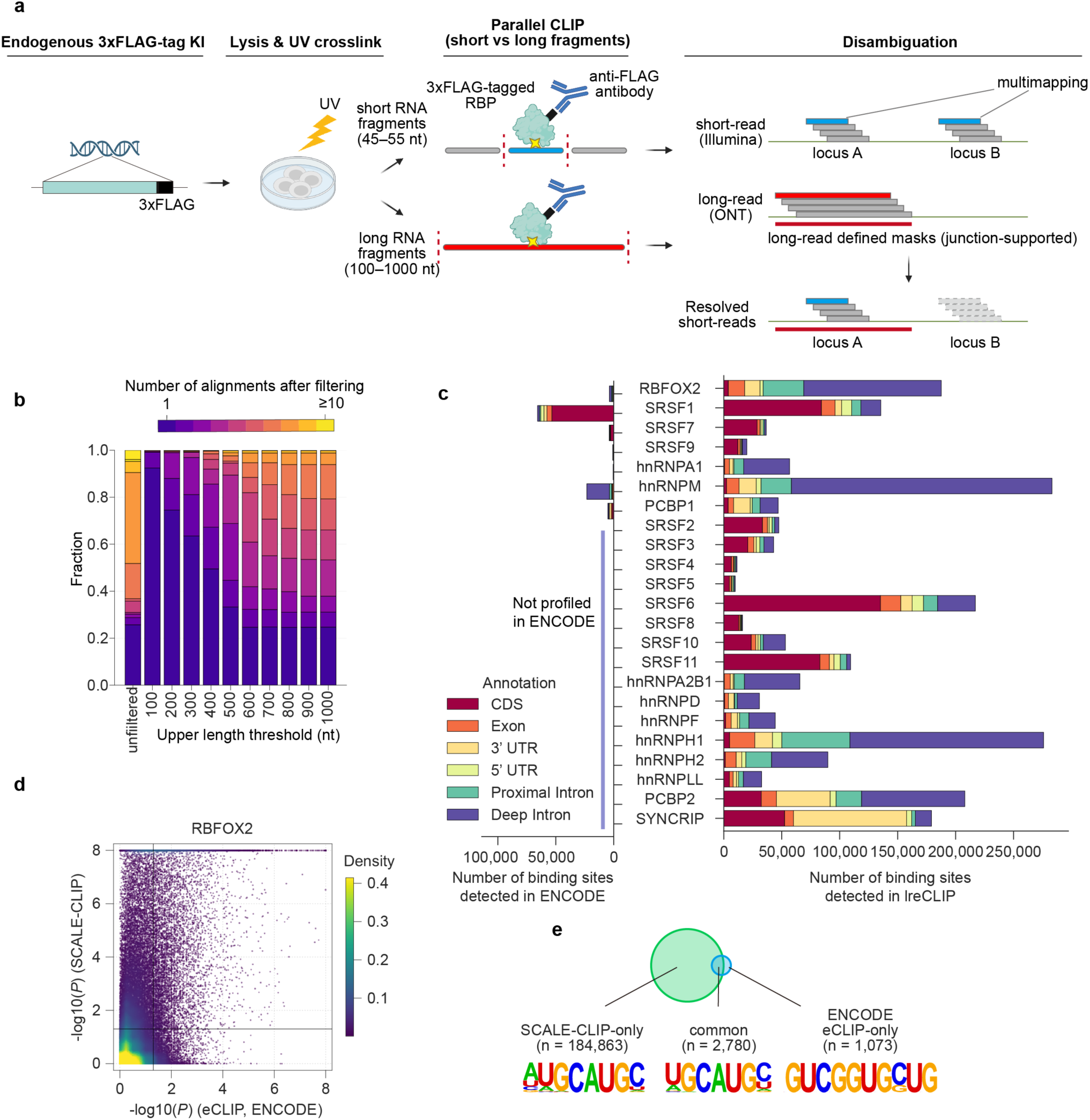
Development and benchmarking of SCALE-CLIP. **a**, Schematic overview of the SCALE-CLIP method. Endogenous RBPs were tagged with 3×FLAG at either the N- or C-terminus using CRISPR-Cas9. Following UV crosslinking, RNA was fragmented into short (45-55 nt) or long (100-1000 nt) fragments. CLIP libraries were sequenced in parallel using short-read (Illumina) and long-read (Nanopore) platforms, and ambiguous or multi-mapped short-read alignments were filtered based on long-read CLIP coverage and overlap. **b**, Stacked bar plots showing the number of candidate alignments per read before and after long-read-guided filtering using mask overlap and splice-junction consistency for each upper length threshold, shown for hnRNPM as a representative example. **c**, Comparison of total number of CLIP peaks and peak distributions across transcript features between ENCODE eCLIP (left) and SCALE-CLIP (right). The top seven SFs are shared between the two datasets, while the remaining 16 SFs are unique to the current dataset (not available in ENCODE K562). **d**, Density-encoded scatter plot comparing window-level CLIP enrichment between ENCODE eCLIP (x-axis) and SCALE-CLIP (y-axis) for seven benchmark SFs. Each point represents a window tested in both datasets; color intensity indicates local point density estimated by fastKDE. Enrichment is defined per window as the maximum −log_10_*P* for IP-versus-input enrichment across replicates. Vertical and horizontal lines indicate unadjusted *P* = 0.05 threshold. **e**, Overlap and motif enrichment analysis of RBFOX2 peak sets classified as SCALE-CLIP-only, ENCODE eCLIP-only or shared. Numbers indicate the number of RBFOX2 peaks in each category. Detailed motif enrichment results are shown in Extended Data Fig. 3f.

We next benchmarked SCALE-CLIP against ENCODE eCLIP to assess peak detection sensitivity, specificity, reproducibility, and RBP coverage under a matched analysis framework (see Methods for details). At matched FDR, SCALE-CLIP substantially increased peak detection across seven shared RBPs, with a median 12.2-fold (2.1–106.7 in range) gain relative to ENCODE eCLIP, while maintaining stringent enrichment and reproducibility (Fig. 1c). In addition, SCALE-CLIP expanded RBP coverage beyond ENCODE by producing robust binding maps for 16 additional RBPs lacking high-quality datasets, with a median of 50,321 (9,900–275,862 in range) peaks per factor (Fig. 1c). Long-read-guided mapping further improved peak detection, increasing the number of supported binding events while removing weakly supported peaks, indicating improved signal recovery with maintained specificity (Extended Data Fig. 1b, c and 3a). Consistent with this, SCALE-CLIP recovered additional peaks in low-uniqueness regions, including known repeat elements (Extended Data Fig. 3b, c). Together, these results demonstrate that SCALE-CLIP improves recovery of endogenous RBP binding events while maintaining specificity and reproducibility. To assess concordance between SCALE-CLIP and ENCODE eCLIP, we compared transcript window-level enrichment scores (Fig. 1d and Extended Data Fig. 3d). For SRSF1, for which the ENCODE dataset showed robust signal, window-level enrichment scores were strongly correlated between datasets, supporting concordant detection of major binding landscapes. For the remaining factors, SCALE-CLIP showed a global shift toward higher enrichment across tiled windows, consistent with increased sensitivity rather than a qualitative change in binding distribution. Replicate reproducibility was comparable between SCALE-CLIP and ENCODE eCLIP, indicating that increased sensitivity does not compromise data consistency (Extended Data Fig. 3e). Sequence-level specificity was also preserved, as SCALE-CLIP peaks showed strong enrichment of canonical RBP motifs, including enhanced enrichment of the established RBFOX2 UGCAUG motif^26^ relative to ENCODE-specific peaks (Fig. 1e and Extended Data Fig. 3f). Together, SCALE-CLIP expands the sensitivity and factor coverage of endogenous RBP mapping while preserving motif specificity and replicate reproducibility, providing a standardized resource for comparing splicing-regulatory RBPs.

### Splicing-regulatory RBPs form structured co-occupancy networks

Leveraging the standardized SCALE-CLIP maps, we next investigated the interaction architecture of splicing-regulatory RBPs. Pairwise comparisons of binding-site overlap resolved three major groups with shared binding preferences, with the most prominent association among SRSF splicing factors (SRSF1-11) (Fig. 2a). Within the SRSF family, strong local co-binding was observed, exemplified by sharp enrichment of SRSF6 signal around SRSF1 binding sites and highly similar exon-centered binding architectures across cassette exons (Fig. 2b and Extended Data Fig. 4a). In contrast, hnRNPs exhibited more heterogeneous and subgroup-specific positional patterns. These group-specific binding architectures are consistent with known differences in protein features and suggest that RBP occupancy is organized into functionally distinct modules rather than uniformly distributed across factors^27^.

**Fig. 2.**
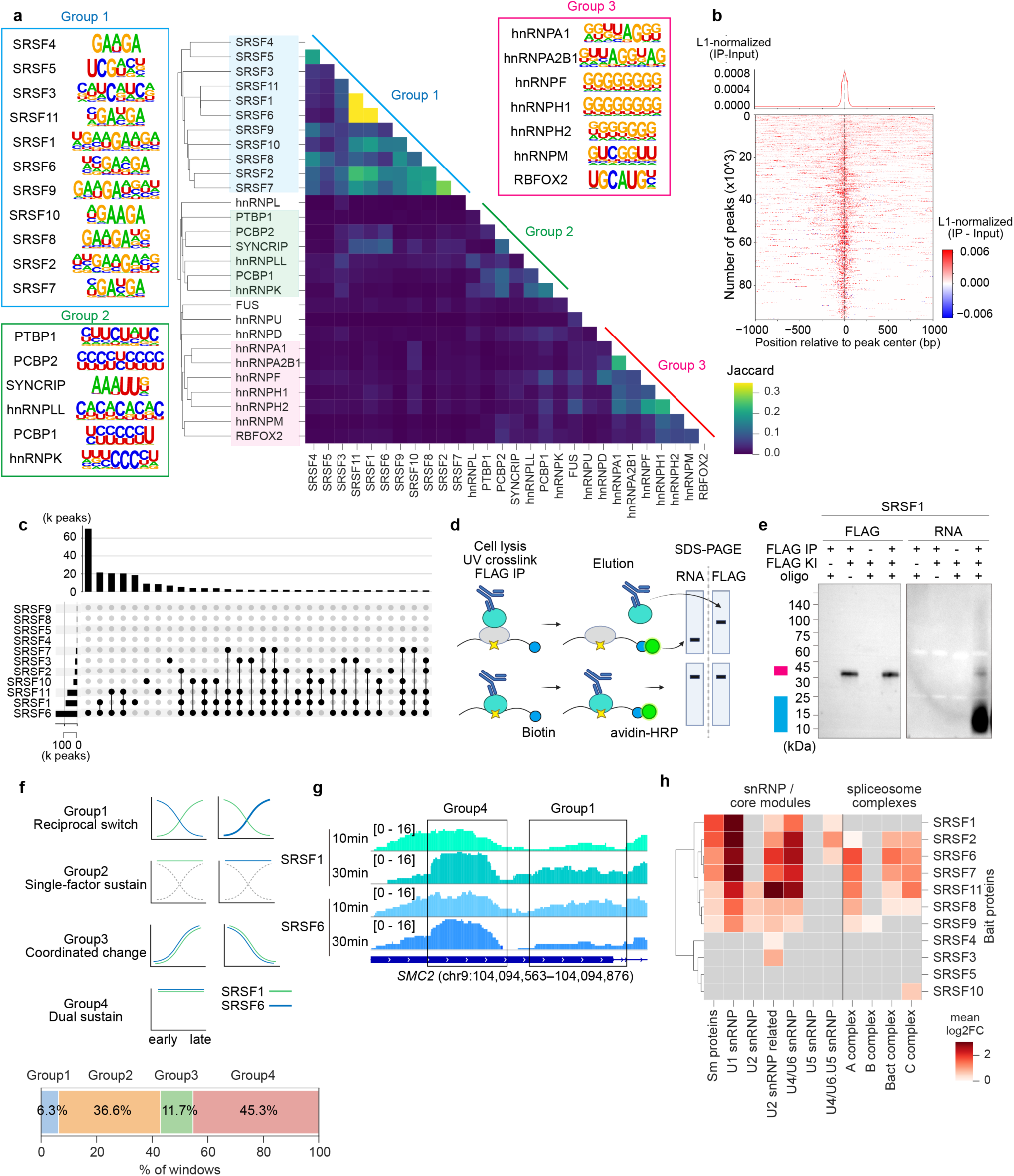
Competitive architecture and temporal dynamics of SRSF co-binding networks. **a,** Pairwise Jaccard indices between reproducible enriched binding windows of RBPs identified by SCALE-CLIP. **b,** Normalized SCALE-CLIP coverage of SRSF6 binding around SRSF1 binding sites (± 1 kb). **c,** UpSet plots showing shared binding peaks among SRSF family proteins. Bars indicate the number of peaks occupied by the indicated factor combinations; only the top 30 intersections by peak count are shown. **d,** Workflow of RNA oligonucleotide binding assay used to distinguish direct RNA binding from indirect association. Biotinylated RNA oligos containing a common SRSF-binding motif (GAAGA) were incubated with cell lysates, UV-crosslinked, immunoprecipitated with anti-FLAG antibody, and detected with avidin-HRP. Direct versus indirect interactions were distinguished based on whether the biotinylated RNA signal co-migrated with the target SF or appeared at distinct molecular weight. **e,** Representative RNA oligonucleotide binding assay results for SRSF1. Magenta bar indicates SRSF1 and the SRSF1-crosslinked biotinylated oligo; cyan bar indicates free biotinylated oligo. Representative result from n = 3 independent experiments with similar results. **f,** Top, schematic of the four temporal binding-transition classes. Bottom, stacked bar plot showing the proportion of windows in each class. **g,** An example genome-browser view of Group1 loci showing reciprocal temporal changes in SRSF1 and SRSF6 binding and Group 4 loci showing dual sustain of SRSF1 and SRSF6 binding. Boxed regions indicate representative intronic intervals. All tracks are shown on the same y-scale. **h,** Enrichment of spliceosome-associated protein classes in SRSF IP-MS datasets. Color indicate median log_2_ fold-enrichment over parental control.

We next quantified higher-order co-occupancy within the SRSF family. Extensive multi-factor binding was observed, with a median of 71% (range, 22–92%) of peaks co-occupied by more than three SRSF proteins (Extended Data Fig. 4b). This co-occupancy was distributed across diverse combinations of SRSF factors, indicating that binding is organized as higher-order assemblies rather than limited to pairwise interactions (Fig. 2c and Extended Data Fig. 4c). To test whether this shared occupancy primarily reflected direct RNA-RBP binding or indirect associations mediated by protein-protein interactions, we UV-crosslinked biotinylated RNA oligonucleotides containing a common SRSF motif to 3×FLAG-tagged RBPs and recovered RBP–RNA complexes by FLAG immunoprecipitation (Fig. 2d). SRSF1, SRSF2 and SRSF7 showed direct binding to the GAAGA motif identified by SCALE-CLIP (Fig. 2e and Extended Data Fig. 4g). These results suggest that at least a subset of multi-SRSF co-occupancy reflects direct RNA binding to shared motif-containing sites. To test whether apparent redundancy within the SRSF network reflects simple site-level displacement, we performed CLIP-seq in SRSF7–3×FLAG cells after SRSF1 knockdown (Extended Data Fig. 4d-f). SRSF1 depletion did not cause a systematic gain of SRSF7 binding at SRSF1-bound sites, arguing against a simple one-to-one displacement model and suggesting that shared occupancy is resolved in a context-dependent manner.

Apparent co-occupancy at the same RNA segment could also reflect temporal exchange of RBPs during RNA processing. Thus we next assessed RBP binding profiles in nascent transcript using 5-ethynyluridine (5-EU) pulse-chase. Time-resolved analysis revealed distinct temporal modes of RBP binding, including reciprocal, persistent and coordinated occupancy patterns between SRSF1 and SRSF6 (Fig. 2f, g). These findings indicate that temporal switching between RBPs contributes to the apparent co-occupancy observed in static SCALE-CLIP profiles. This temporal organization may reflect sequential engagement of SRSF family members at different stages of co-transcriptional splicing and RNA maturation. To examine whether distinct occupancy behaviors are linked to spliceosome assembly, we profiled SRSF-associated proteomes by IP-MS (Fig. 2h and Extended Data Fig. 4h, i). Notably, SRSF-associated proteomes showed diverse enrichment of snRNP components and spliceosomal subcomplexes, consistent with engagement at distinct stages of spliceosome assembly (Fig. 2h and Supplementary Data 4). Together, these analyses suggest that shared SRSF occupancy represents a mixture of direct motif recognition, temporal exchange and spliceosome-associated states, providing a mechanistic context for the positional splicing rules defined below.

### SRSF binding position determines splicing regulatory direction

To define how RBP binding position relates to splicing outcomes, we integrated SCALE-CLIP binding profiles with alternative splicing changes following RBP knockdown (Extended Data Fig. 5a–d). Long-read transcript assembly further expanded detection of perturbation-induced isoforms, capturing additional splicing events beyond reference annotations (Extended Data Fig. 5c).

We next constructed splicing maps to quantify position-dependent binding effects across knockdown-responsive events (Fig. 3a)^28^. Several factors including RBFOX2, hnRNPM and hnRNPH1 showed positional patterns consistent with previous reports, in which binding in flanking introns was associated with alternative exon inclusion or skipping^29–32^(Fig. 3a and Extended Data Fig. 5e). Among the factors examined, SRSF family proteins showed the most consistent family-wide position-dependent associations with splicing outcomes. Consistent with canonical models^33^, binding of SRSF family proteins within alternative exons was associated with exon inclusion. Strikingly, exons that became more included after SRSF knockdown showed enriched SRSF binding on upstream or downstream flanking exons, indicating that flanking-exon SRSF occupancy acts as a skipping-promoting input (Fig. 3a, b). These opposite positional associations suggest that the SRSF regulatory output is determined primarily by binding location, establishing a positional rule in which alternative-exon and flanking-exon occupancy exert opposite effects on exon inclusion.

**Fig. 3.**
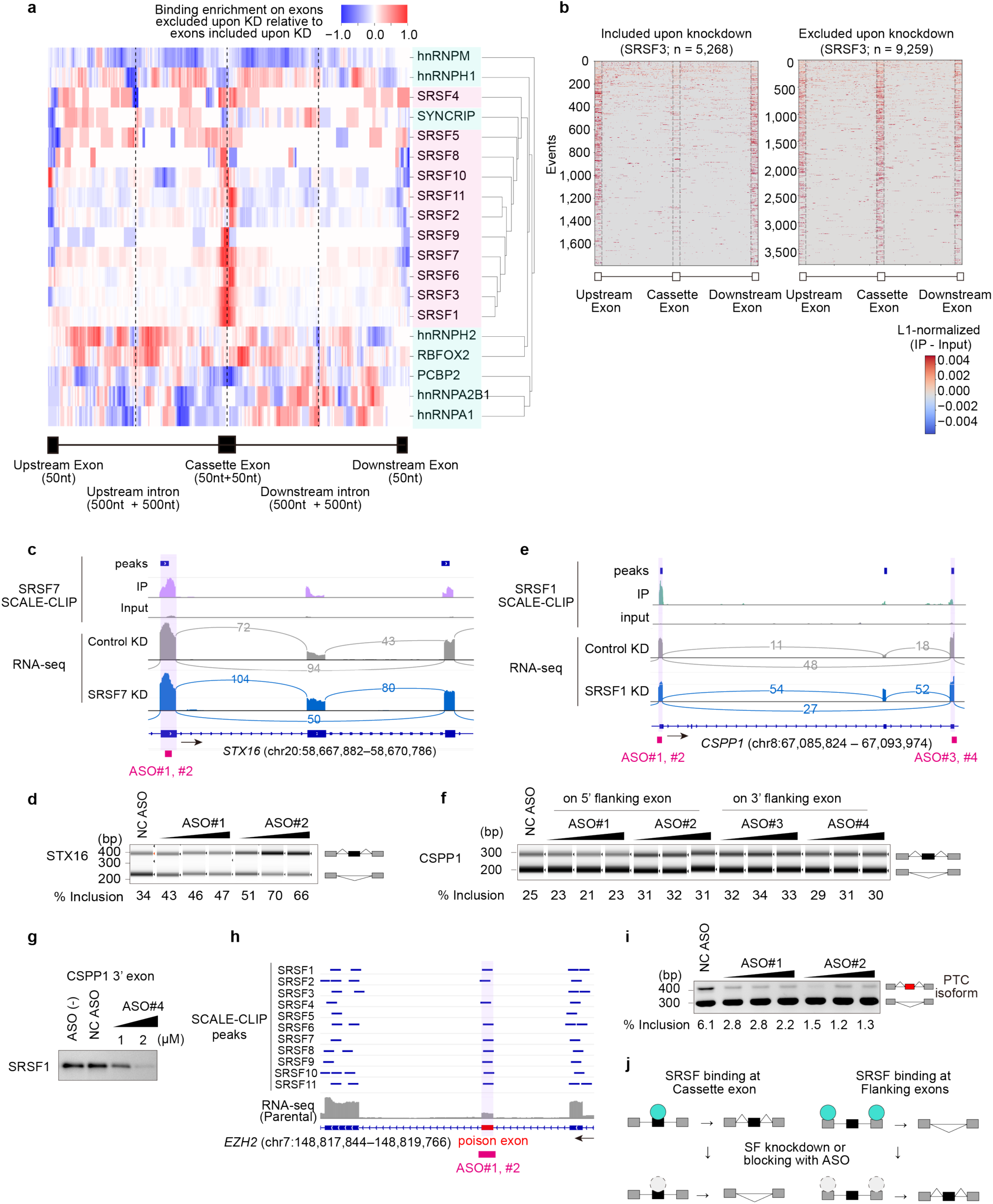
Integration of SCALE-CLIP peaks with splicing changes reveals bidirectional splicing regulation. **a,** Differential SCALE-CLIP peak-density profiles around cassette exons that are excluded or included upon knockdown (KD) (KD RNA-seq; |ΔPSI| > 5% and FDR < 0.05). Δbinding was defined as density(excluded) – density(included). RBPs were clustered by similarity of Δbinding profiles. **b,** SRSF3 SCALE-CLIP signal around cassette exons included upon KD (left) or skipped upon KD (right). **c, d,** Representative loci for SRSF7 (c) and SRSF1 (d), showing SRSF binding associated with increased cassette exon inclusion upon KD. Upper tracks, SCALE-CLIP peaks; lower tracks, KD and control RNA-seq coverage. Magenta bars, ASO target regions. **e, f,** RT–PCR validation of STX16 (e) and CSPP1 (f) splicing after transfection of two independent ASOs targeting SCALE-CLIP-defined SRSF binding sites (0.05, 0.1 or 0.3 µM, 24 h) or a non-targeting control ASO. **g,** RNA oligo pull-down followed by anti-FLAG immunoblot showing dose-dependent binding of SRSF1-3×FLAG to an RNA oligo from the CSPP1 downstream flanking exon, with targeting ASO (1 or 2 µM) or non-targeting control ASO. **h,** SRSF family proteins bind the EZH2 poison exon. Two ASOs were designed on this region (magenta bars). **i,** RT-PCR validation of EZH2 splicing after transfection of two independent ASOs targeting SCALE-CLIP-defined SRSF binding sites (0.05, 0.1 or 0.3 µM, 24 h) or a non-targeting control ASO. Cells were treated with cycloheximide for 5 h before harvest. **j,** Model summarizing the positional rules linking SRSF binding topology to splicing outcomes.

We next tested whether flanking-exon SRSF occupancy causally suppresses cassette exon inclusion using antisense oligonucleotides (ASOs) designed to block SCALE-CLIP-defined binding sites. ASO-mediated masking of flanking-exon SRSF sites increased cassette exon inclusion at multiple loci in a dose-dependent manner, supporting a causal role for flanking-exon occupancy as a repressive input (Fig. 3c-f and Extended Data Fig. 6a-d). ASO efficacy was confirmed by reduced SRSF binding at targeted sites (Fig. 3g). At the well-established SRSF2 target EZH2 poison exon^34^, ASO-mediated blockade of multiple SRSF binding reduced exon inclusion, consistent with the canonical inclusion-promoting role of exonic SRSFs (Fig. 3h, i and Extended Data Fig. 6e, f). This positional contrast demonstrates that regulatory direction is not dictated by motif identity alone: masking similar SRSF-recognition motifs produced opposite splicing outcomes depending on whether the sites resided in alternative or flanking exons. Together, these results establish a positional SRSF-binding rule in which the same class of RNA motifs can promote inclusion or skipping depending on exon context (Fig. 3j).

### Combinatorial SRSF occupancy tunes the positional splicing code

We next asked how higher-order SRSF occupancy modulates this positional rule and influences basal exon inclusion and perturbation responsiveness. Focusing on alternative exons with suboptimal splice sites (Methods and Extended Data Fig. 7a), multi-member SRSF occupancy within alternative exons was associated with higher baseline inclusion and reduced sensitivity to knockdown, compared with binding by the perturbed SRSF alone (Fig. 4a, b).

**Fig. 4.**
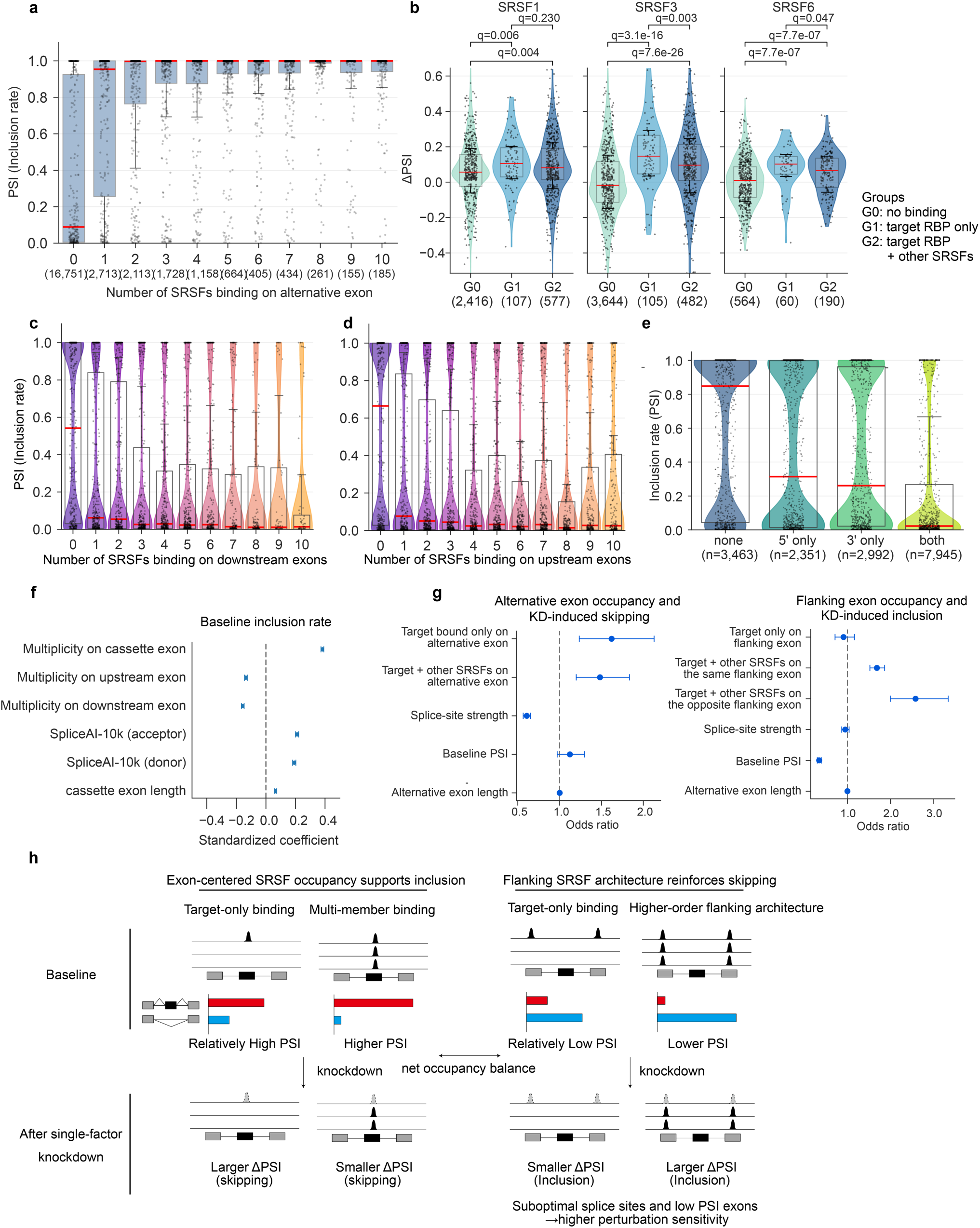
Multi-member SRSF organization tunes exon state and perturbation sensitivity. **a,** PSI distributions for alternative exons with suboptimal splice sites, stratified by SRSF binding multiplicity on the alternative exon. **b,** ΔPSI distribution for significantly changed events (FDR < 0.05) after SRSF1, SRSF3, or SRSF6 knockdown, restricted to events with suboptimal splice sites (SpliceAI-10k < 0.5), stratified by SRSF binding at the cassette exon: no SRSF binding, binding by the knocked-down SRSF only, or co-binding by the knocked-down SRSF and ≥1 additional SRSF family member. Positive ΔPSI indicates increased exon inclusion upon knockdown. **c–e,** PSI distributions for alternative exons with suboptimal splice sites and no detectable SRSF binding on the alternative exon, plotted by SRSF binding multiplicity on the upstream (**c**) or downstream (**d**) flanking exon, or by flanking-exon binding pattern (upstream only, downstream only, or both; **e**). Additional strata are shown in Extended Data Fig. 7c–e. For **a** and **c–e**, suboptimal splice sites were defined as SpliceAI-10k < 0.6 for both donor and acceptor sites. Points represent individual exons. Boxplots show the median, interquartile range and 1.5 × IQR. Group sizes are shown as the number of alternative exons in **a** and **e**, and significant splicing events in **b**. **f,** Multivariable linear regression model of baseline exon inclusion. Points indicate standardized regression coefficients and horizontal bars indicate 95% confidence intervals. **g,** Logistic regression models of directional perturbation sensitivity, showing features associated with knockdown-induced skipping (left) and inclusion (right). Points indicate odds ratios and horizontal bars indicate 95% confidence intervals. **h,** Model summarizing exon-centered buffering, flanking repression, and perturbation-sensitive exon architecture.

Stratifying exons by SRSF multiplicity revealed that co-occupancy amplifies the opposing effects of alternative-exon and flanking-exon binding. Increasing SRSF occupancy within alternative exons remained positively associated with exon inclusion, whereas increasing multiplicity on either upstream or downstream flanking exons was associated with progressively lower inclusion (Fig. 4c, d and Extended Data Fig. 7b–d). These results establish alternative-exon and flanking-exon SRSF binding as opposing positional inputs, whose balance collectively contributes to steady-state exon inclusion levels.

Direct comparison of flanking configurations further revealed an asymmetry, with downstream flanking occupancy exerting stronger repressive effects than upstream binding, and simultaneous occupancy on both flanking exons producing the lowest inclusion levels (Fig. 4e and Extended Data Fig. 7e). To quantitatively model these effects, we fitted multivariate linear regression models of baseline exon inclusion. Alternative-exon SRSF multiplicity and splice-site strength were positively associated with inclusion, whereas SRSF multiplicity on upstream or downstream flanking exons contributed negatively, consistent with opposing positional effects (Fig. 4f). These results formalize SRSF occupancy as asymmetric and opposing positional inputs that explain exon inclusion beyond splice-site strength and exon length. A covariates-only model including splice-site strength and cassette exon length explained 21.0% of the variance in baseline PSI (R² = 0.210). Incorporating positional SRSF multiplicity increased the explained variance to 32.2% (R² = 0.322), outperforming reduced models containing either cassette-exon SRSF multiplicity alone (27.4%, R² = 0.274) or flanking-exon SRSF multiplicity alone (21.5%, R² = 0.215). These results indicate that the combined positional architecture captures regulatory information that is not explained by splice-site strength, exon length, or single-position SRSF occupancy alone.

We next examined how exonic binding architecture influences sensitivity to SRSF perturbation. Suboptimal splice sites were associated with increased responsiveness to knockdown, affecting both exon skipping for alternative exon-bound events and exon inclusion for flanking-bound events (Fig. 4g). Among alternative exon-bound events, target-only binding showed greater sensitivity than multi-member binding, consistent with buffering by exon-centered co-occupancy. In contrast, among flanking-bound exons, lower baseline inclusion and higher-order flanking occupancy were associated with increased sensitivity to perturbation, suggesting that multiple SRSF occupancy on flanking exons marks a perturbation-sensitive repressive architecture (Fig. 4g). Together, these results indicate that combinatorial SRSF organization tunes both steady-state exon inclusion and sensitivity to perturbation: exon-centred co-occupancy stabilizes inclusion, whereas flanking-exon architectures reinforce repression and increase responsiveness (Fig. 4h).

### RNA modifications are coupled to RBP binding landscapes

RNA modifications are increasingly recognized as regulators of RNA processing^35^, yet their impact on RBP binding landscapes remains poorly characterized. We profiled epitranscriptomic marks on nascent transcripts in chromosome-associated RNA (caRNA) and integrated these data with SCALE-CLIP binding maps (Extended Data Fig. 9a-c and Methods). We observed broad associations between RBP binding and RNA modification marks, with the strongest enrichment occurring between SRSF peaks and highly modified m^6^A sites (Fig. 5a–c and Extended Data Fig. 8b–e). Multiple SRSFs were enriched at m^6^A sites, and m^6^A-overlapping SRSF peaks were predominantly exonic, indicating preferential coupling between m^6^A-marked exonic regions and SRSF occupancy (Fig. 5c, d). Beyond m^6^A, PTBP1 showed preferential binding at Ψ-rich sites, and several SRSFs exhibited enrichment at m^5^C sites (Extended Data Fig. 8b–e). Perturbing m^6^A deposition altered SRSF binding at a subset of loci, supporting a functional link between RNA modification state and RBP occupancy (Extended Data Fig. 9d–f). Together, these results indicate that RNA modifications modulate RBP binding landscapes, providing an additional regulatory layer for splicing-associated RBP occupancy.

**Fig. 5.**
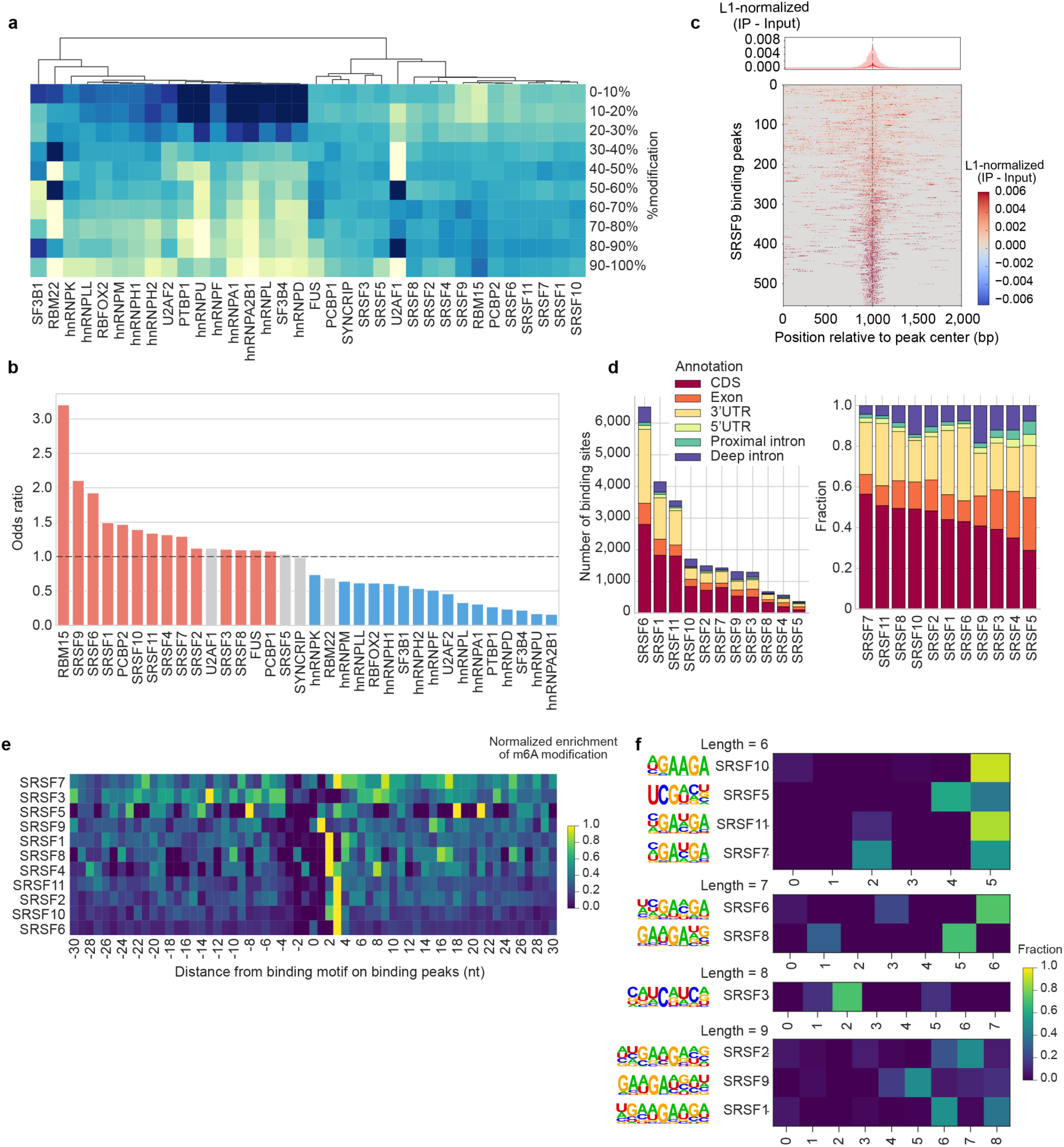
Epitranscriptomic signatures in splicing factor binding maps. **a,** Distribution of CLIP-m^6^A overlap across m^6^A modification-frequency bins (0-100%, 10% increments) for each RBP. For each bin, the overlap rate was calculated as the fraction of m^6^A-called bases within an RBP peak and normalized within each RBP. RBPs were hierarchically clustered. **b,** Enrichment of RBP binding at highly modified m^6^A sites. Odds ratios compare peak overlap at highly modified (≥ 30%) versus low-modification (≤ 30%) sites. Significance was assessed by two-sided Fisher’s exact test with Benjamini-Hochberg correction (FDR < 0.05); red and blue indicate significant enrichment and depletion, respectively; grey, not significant. **c,** Metaprofile of L1-normalized SRSF9 SCALE-CLIP coverage centered on highly-modified m^6^A bases (≥ 80%, ± 1 kb). **d,** Transcript-feature annotation of SRSF binding peaks overlapping m^6^A sites (≥ 30%), shown as counts and fractions. **e,** Motif-centered metaprofile of high-confidence m^6^A sites (≥ 30%) within RBP binding peaks, anchored on the top HOMER motif for each RBP. Profiles were computed in a strand-aware manner and normalized to a maximum of 1. **f,** Representative example for SRSF4, shown as in (**e**). **g,** Distribution of high-confidence m^6^A sites within motif bodies. Values are normalized to a maximum of 1. Motifs are grouped by length, and the corresponding top-enriched motifs are shown.

To determine the spatial relationship between m^6^A and SRSF binding motifs, we aligned SCALE-CLIP peaks by motif instances and quantified base-level m^6^A distributions (Fig. 5e, f and Extended Data Fig. 9g). m^6^A was preferentially positioned at the terminal adenosine within the GAAGA motif, indicating a consistent spatial coupling between modification sites and SRSF recognition elements (Fig. 5f). This precise alignment suggests that m^6^A can locally tune SRSF binding at motif-containing sites, providing a potential mechanistic basis for the enrichment of SRSF binding at modified sites. Collectively, these data identify an epitranscriptomic layer in which m^6^A is spatially coupled to SRSF motifs and can modulate local RBP occupancy.

### m^6^A potentiates SRSF-dependent exon inclusion

To test whether the modification-dependent changes in SRSF affinity translate into splicing outcomes, we integrated splice-site strength, base-resolved RNA modification maps, and SCALE-CLIP binding profiles. Exons with higher m^6^A modification levels exhibited higher steady-state inclusion (Fig. 6a). We then stratified exons by the presence of SRSF binding and m^6^A modification within the alternative exon. Exons harboring both SRSF binding and m^6^A showed higher inclusion than those with either feature, consistent with cooperation between modification state and SRSF occupancy (Fig. 6b). These results support a model in which m^6^A creates an epitranscriptomic context that potentiates the inclusion-promoting activity of exon-bound SRSFs.

**Fig. 6.**
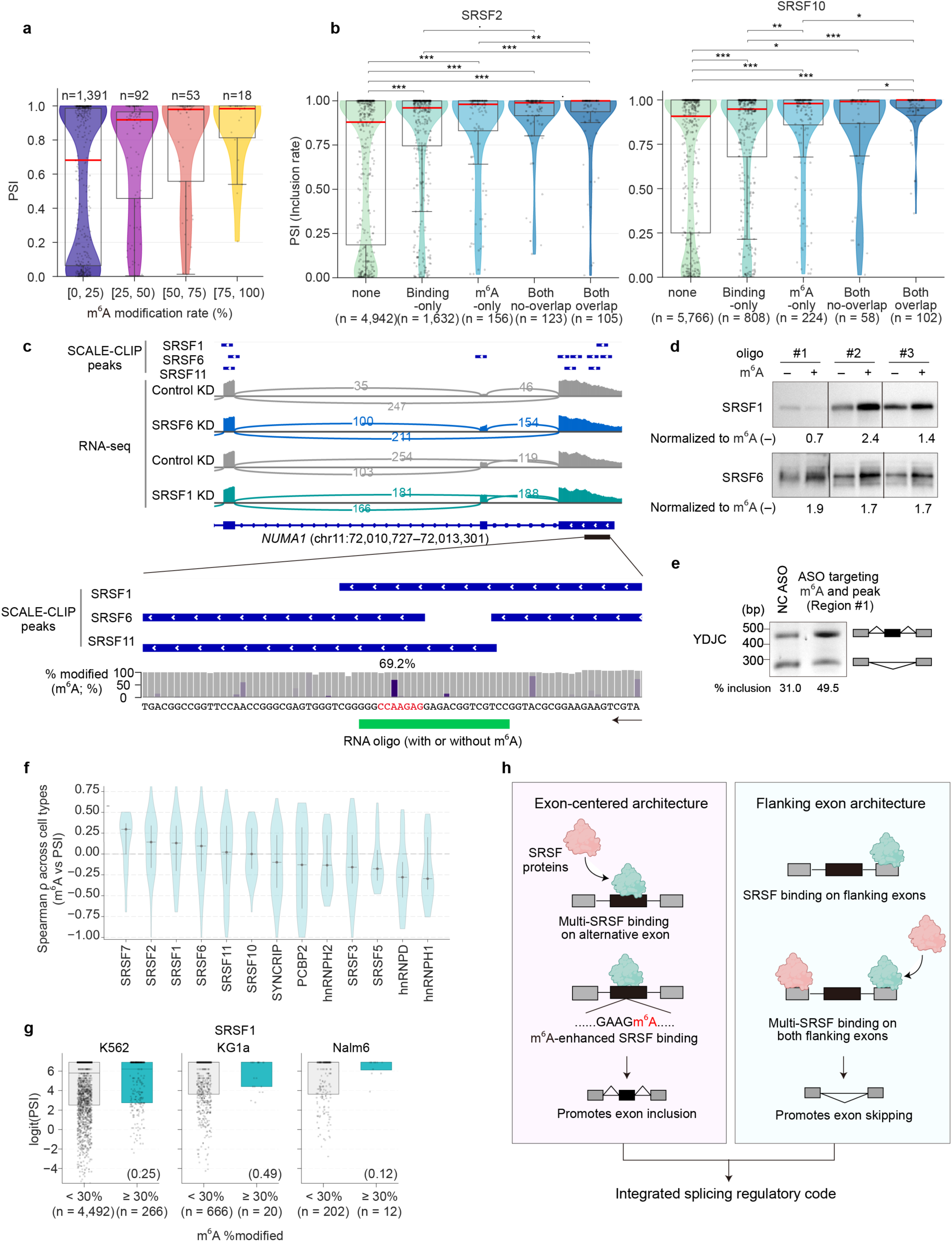
Epitranscriptomic layer of splicing regulation via SRSF binding modulation. **a,** Relationship between exon inclusion (PSI) and m^6^A modification in K562 cells. Percent m^6^A was quantified at m^6^A-called adenosines within cassette exons from caRNA direct RNA-seq, and PSI was obtained from matched RNA-seq. **b,** PSI distributions of alternative exons stratified by SRSF binding and m^6^A states within the alternative exon: none, SRSF-binding only, m^6^A only, both without overlap, and both with overlap. Brackets indicate selected pairwise group comparisons. *P* values were calculated by two-sided Dunn’s test with Benjamini-Hochberg correction; ・*P* < 0.1, \**P* < 0.05, \*\**P* < 0.01, \*\*\**P* < 0.001. For **a** and **b**, Points represent individual cassette exons. Boxplots show the median, interquartile range and 1.5 × IQR. Group sizes indicate the number of cassette exons. **c,** Genome-browser view showing a SCALE-CLIP-defined SRSF peak, the corresponding knockdown-associated splicing change, and base-resolved m^6^A calls. The region used for oligo #3 is highlighted. **d,** Biotinylated RNA oligos (20–30 nt) were synthesized as matched A and m^6^A pairs and incubated with lysates expressing 3×FLAG-tagged SRSF1 and SRSF6, followed by streptavidin pull-down and anti-FLAG immunoblot. A representative experiment is shown. Band intensities should be compared within each matched A/m^6^A pair. **e,** RT-PCR validation after transfection of an ASO targeting a region overlapping the predicted m^6^A site and the SRSF6 peak at the indicated locus, compared with a non-targeting control ASO. %inclusion was calculated from PCR product length-corrected band intensities. **f,** Distributions of cross-cell-type Spearman correlations between m^6^A level and exon inclusion for RBP-overlapping m^6^A site-event pairs. **g,** Cell-type-specific PSI distributions for cassette exons linked to SRSF1-overlapping m^6^A sites, stratified by m^6^A level at the corresponding site (< 30% or ≥ 30%). Points represent individual cassette exon–m^6^A site pairs. boxplots summarize the proposed relationship among m^6^A, effective SRSF occupancy and splicing outcomes. Boxplots show the median, interquartile range and 1.5 × IQR. Group sizes indicate the number of cassette exon–m^6^A site pairs. *P* values from two-sided Mann-Whitney U tests are shown in parentheses. **h,** Model summarizing the integrated splicing regulatory code defined by positional, combinatorial and epitranscriptomic layers.

To test whether m^6^A directly modulates SRSF-RNA affinity, we examined SRSF binding to matched RNA substrates differing only at a single modified nucleotide within endogenous SCALE-CLIP sites (Fig. 6c). m^6^A altered SRSF binding across multiple loci, as measured by RNA pull-down assays with SRSF1, SRSF6, and SRSF11, although the magnitude and direction of the effect varied by sequence context (Fig. 6d and Extended Data Fig. 10a, b). Consistent with functional relevance, targeting the m^6^A-containing SRSF6 binding element at the YDJC locus with an ASO modulated splicing in the predicted direction, supporting its role as an endogenous regulatory site (Fig. 6e). These results demonstrate that m^6^A can directly and locally enhance SRSF binding at splicing-associated sites, providing a mechanistic basis for the observed coupling between RNA modification and splicing regulation.

Finally, we examined whether the observed m^6^A-SRSF-splicing relationship extends across diverse cell types. Analysis of RNA modification and splicing profiles across multiple human cell types revealed concordant m^6^A levels at SRSF-associated sites, indicating that a subset of these modification contexts is broadly conserved (Extended Data Fig. 10c, d). At the level of splicing output, m^6^A at SRSF-bound sites was consistently associated with increased exon inclusion across cell types, as observed for multiple SRSFs including SRSF7, SRSF2, SRSF1, and SRSF6 (Fig. 6f, g and Extended Data Fig. 10e–g). These findings indicate that m^6^A-dependent tuning of SRSF binding is linked to exon inclusion across diverse cellular contexts. Together, our results support a multi-layered model in which positional SRSF binding, higher-order occupancy and m^6^A modification jointly shape exon inclusion outcomes (Fig. 6h).

## Discussion

Here we present SCALE-CLIP, an endogenous and cross-comparable framework for transcriptome-wide mapping of RBP binding, and combine it with epitranscriptomic profiling to define a multi-layered RNA-binding code for alternative splicing. By integrating CRISPR-Cas9-mediated endogenous epitope tagging with long-read-guided assignment of short-read CLIP reads, SCALE-CLIP generates directly comparable binding maps across splicing-regulatory factors while improving recovery of binding events in repeat-rich and multimapping regions. This framework reveals how binding position, multi-factor organization and RNA modification context are integrated to shape exon choice at the transcriptome scale.

A central technical advance of SCALE-CLIP is LRA, which mitigates the multimapping problem of conventional short-read CLIP in repeat-rich and low-uniqueness regions^36,37^. Coupled to endogenous epitope tagging, this strategy enables systematic and directly comparable mapping across RBPs under physiological conditions, reducing dependency on antibody quality and avoiding perturbative overexpression systems^16–18^. Extending SCALE-CLIP across major splicing-regulatory RBPs establishes that splicing regulation emerges from structured, multi-factor interaction networks, rather than isolated single-factor occupancy. These networks encompass co-occupancy, competition and dynamic exchange among RBPs, providing a more integrated view of how splicing decisions are encoded at the transcriptome scale. While prior studies have provided individual examples of SRSF cooperation and competition, direct family-scale comparisons have been constrained by overexpression, endogenous-antibody variability, and limited factor coverage^18,38,39^. By combining SCALE-CLIP with UV-crosslinked RBP immunoprecipitation and time-resolved profiling, we show that shared binding sites resolve into temporally distinct occupancy states, including mutually exclusive, factor-primed and co-enriched configurations. These findings suggest that apparent redundancy among SRSFs reflects a temporally organized and dynamic regulatory system, rather than simple static co-binding at shared RNA elements.

Integration of SCALE-CLIP with splicing analyses revealed a positional code for SRSF-dependent splicing regulation, in which binding within alternative exons promotes exon inclusion, whereas binding on flanking exons promotes skipping. This extends the RNA splicing-map framework^40^ established for Nova, PTB and Rbfox proteins^41–43^ to directly comparable endogenous binding maps across the SRSF family at transcriptome scale. These position-dependent effects are further modulated by combinatorial SRSF organization in an asymmetric manner. On alternative exons, higher-order co-occupancy stabilizes inclusion and buffers sensitivity to perturbation, providing a mechanistic basis for splicing robustness. In contrast, on flanking exons, broader SRSF occupancy reinforces repression and increases responsiveness to perturbation. This inversion suggests that regulatory outcome is not specified by individual binding alone, but by higher-order positional architectures of SRSF occupancy. The increased perturbation sensitivity associated with bilateral flanking-exon occupancy further suggests that such architectures may coordinate repressive inputs from both flanking positions, thereby reinforcing exon skipping through bilateral occupancy. We further provide proof-of-principle that the splicing code defined here is mechanistically actionable, by targeting motif-enriched SCALE-CLIP peaks with splice-switching ASOs and shifting exon inclusion in the predicted direction across multiple loci. Notably, exons embedded within broader flanking SRSF architectures, as well as those with weaker splice sites and lower baseline inclusion, showed greater responsiveness to perturbation, suggesting that these features can serve as practical criteria for prioritizing SCALE-CLIP-guided ASO target selection. These results indicate that map-guided ASO design can narrow the empirical search space relative to conventional ASO walking approaches and may enable more rational modulation of splicing outcomes, highlighting the potential for rational, code-informed splicing modulation^44,45^.

Our findings reveal that m^6^A constitutes an epitranscriptomic layer that can locally tune SRSF binding affinity and thereby influence splicing outcomes, extending the repertoire of m^6^A-responsive splicing regulation beyond core spliceosomal components to canonical auxiliary factors. This coupling is most evident at m^6^A-marked SRSF binding sites, where modification state is associated with increased SRSF occupancy and higher exon inclusion. Beyond these associations, matched RNA pull-down assays provide orthogonal support for a mechanistic link, showing that m^6^A at the same SCALE-CLIP-defined sequence can locally alter SRSF binding, consistent with cis-proximal, sequence-dependent tuning of binding potential. RNA modifications have been shown to directly affect splice-site recognition by core spliceosomal factors, including m6A-mediated inhibition of U2AF35 binding at the 3′ splice site and pseudouridine-dependent weakening of U2AF65 recognition in polypyrimidine tracts^7,8,46^. Our findings extend this concept to auxiliary splicing factors and complement established routes by which m^6^A reshapes RBP binding, including reader and writer-mediated recruitment/antagonism^9,47^ and structure-dependent access control^48^.

Several limitations should be considered. Peak recovery remains influenced by transcript abundance and long-read coverage, particularly at low-expression loci, and tag- or immunocapture-specific effects cannot be fully excluded. In addition, the current framework uses reference-genome-based analyses and does not explicitly model genetic variation in cis-regulatory elements or trans-acting factors. Incorporating deeper long-read coverage, orthogonal validation and individual genetic variation should extend SCALE-CLIP from general regulatory principles toward disease- and patient-specific models of splicing regulation.

The standardized binding maps, positional rules, and epitranscriptomic integration defined here establish a quantitative framework for interpreting splicing regulation. By integrating RBP occupancy with motif, positional, and modification features, this framework provides a basis for estimating how sequence variation, including disease-associated single-nucleotide variants, may perturb RBP binding and alter splicing at specific loci. More broadly, this work advances RBP binding maps from descriptive resources toward quantitative models of regulatory output, providing a foundation for mechanistic interrogation, rational splicing modulation and interpretation of splicing dysregulation in human disease.

## Materials and Methods

### Cell lines

K562, Nalm-6 and KG-1a cells were cultured in RPMI1640 medium supplemented with 10% fetal bovine serum and 100 U/ml penicillin/streptomycin; A549, HEL, HeLa and HepG2 cells were cultured in DMEM medium supplemented with 10% fetal bovine serum and 100 U/ml penicillin/streptomycin. All cells were maintained at 37 °C in a humidified 5% CO₂ atmosphere. Cells were routinely tested for mycoplasma contamination with PCR Mycoplasma Detection Set (TaKaRa).

### Genome editing with CRISPR-Cas9

Guide RNAs (gRNAs) and single-stranded oligodeoxynucleotide (ssODN) donor templates to insert 3×FLAG-tag to either N- or C-terminus were designed and synthesized by Integrated DNA Technologies (IDT). Tag insertion sites were chosen to maximize distance from the RRM (RNA recognition motif) while prioritizing sites with favorable cut-to-insert distances and predicted on-/off-target scores. The gRNA, donor template, and 3×FLAG insertion site (N or C-terminal) are provided in Supplementary Data 6 for each SF. For ribonucleoprotein (RNP) complex assembly, synthetic crRNA and tracrRNA (IDT) were annealed and incubated with Alt-R S.p. Cas9-GFP V3 protein (IDT). The preassembled RNP and donor ssODN were introduced into parental K562 cells using the Neon NxT Electroporation System (Invitrogen) following the manufacturer’s protocol. GFP-positive cells were single-cell sorted into 96-well plates using a BD FACSAriaIII and expanded for clonal isolation. Correct in-frame insertion of the 3×FLAG tag at the target locus was confirmed by genomic PCR, Sanger sequencing and Western blotting with anti-FLAG M2 antibody (Sigma-Aldrich, F1804), followed by Western blotting after siRNA-mediated knockdown of the target gene.

### Long-read-guided enhanced CLIP-seq

#### Short-read seCLIP library preparation

Short-read seCLIP (single-end enhanced CLIP) experiments were performed as previously described^49^, with minor modifications. Briefly, 20 million cells were UV-crosslinked on ice (254 nm, 400 mJ/cm^2^), lysed in buffer containing 50 mM Tris-HCl (pH 7.4), 100 mM NaCl, 1% (vol/vol) Igepal CA-630, 0.1% (vol/vol) SDS and 0.5% (wt/vol) sodium deoxycholate, and sonicated. RNA was fragmented by RNase I treatment to an average size of 45–55 nt. Lysates were incubated with anti-FLAG M2 magnetic beads (Millipore, M8823) for 2–3 h at 4 °C. Two percent (vol/vol) of each lysate was reserved as a size-matched input control. After extensive high- and low-salt washes, RNA-protein complexes were treated with FastAP and T4 PNK (NEB), followed by ligation of a 3′ RNA adapter using T4 RNA ligase I (NEB). High-salt wash buffer contained 50 mM Tris-HCl (pH 7.4), 1 M NaCl, 1% (vol/vol) Igepal CA-630, 1 mM EDTA, 0.1% (vol/vol) SDS and 0.5% (wt/vol) sodium deoxycholate. Low-salt wash buffer contained 20 mM Tris-HCl (pH 7.4), 10 mM MgCl_2_, 0.2% (vol/vol) Tween-20 and 5 mM NaCl. IP efficiency was validated by western blot using DYKDDDDK tag Polyclonal antibody (Proteintech, 20543-1-AP). Pulled-down RNA was visualized by pCp-biotin ligation (Jena Bioscience, NU-1706-BIO) followed by chemiluminescent detection (Thermo Scientific, 89880). IP and input samples were resolved by SDS-PAGE and transferred to nitrocellulose membranes. Membrane regions spanning from the expected protein size to ∼75 kDa above were excised, treated with proteinase K, and RNA was recovered, reverse transcribed (SuperScript IV, Invitrogen), ligated to a 3′ DNA adapter, PCR amplified (Q5 polymerase, NEB), and size selected by gel electrophoresis. Libraries were sequenced on an Illumina NextSeq instrument, with ≥ 20 million reads per replicate. Each seCLIP experiment consisted of IP from three independent biosamples, with a size-matched input for each replicate.

#### Long-read CLIP library preparation

Approximately 20 million cells were UV crosslinked on ice at 400 mJ/cm^2^ with 254 nm radiation and lysed, followed by sonication to generate RNA fragments predominantly in the ∼100-1000 nt range (assessed with Tapestation). Lysates were incubated with anti-FLAG M2 magnetic beads (Millipore, M8823) for 2-3 h at 4 °C. After extensive high- and low-salt washes, RNA-protein complexes were digested with proteinase K, and RNA was recovered using RNA Clean & Concentrator-5 kit (Zymo Research). Ribosomal RNA was depleted with RiboMinus Eukaryote Kit v2 (Invitrogen, A15020) and the remaining RNA was polyadenylated with poly(A) polymerase (Takara, 2180A). cDNA libraries were prepared using the PCR-cDNA Sequencing Kit (SQK-PCS114, Oxford Nanopore Technologies) according to the manufacturer’s protocol and sequenced on a PromethION instrument with Flow Cell R10 (FLO-PRO114M, ONT).

### Primary processing of short- and long-read data

For short-read CLIP reads, adapters were trimmed with skewer (v0.2.2)^50^ and UMIs (10 nt) were extracted and appended to read names using fastp (v0.24.0)^51^ for downstream deduplication. Reads were aligned to reference human genome (GRCh38, no-alt-analysis set) with GENCODE v44 annotation using STAR (v2.7.10b)^52^, retaining multimappers for subsequent disambiguation (--outFilterMultimapNmax 100 --winAnchorMultimapNmax 100 --outSAMmultNmax 100 --outSAMtype BAM SortedByCoordinate). For long-read CLIP reads, raw pod5 files were basecalled with Dorado version 1.0.1 (ONT) in high-accuracy mode. Reads were subsequently filtered (mean Q score ≥ 9, read length ≥ 50 nt) and oriented using Pychopper (v2.7.10), and only full-length oriented reads were aligned to the same reference genome using minimap2 (v2.28) in splice-aware mode (-ax splice -uf -k14 --secondary=no)^53^. Alignments were filtered to retain primary alignments with MAPQ ≥ 20.

### Long-read-guided read attribution (LRA) of short-read multimappers

Uniquely mapped short reads were retained as aligned. For multimapping short reads, candidate alignments were filtered using long-read evidence to enforce exon- and junction-level concordance (workflow in **Extended Data Fig.** 1a). Long-read exon blocks were derived from long-read CLIP alignments. For each candidate short-read placement, strand-consistent overlap with long-read exon blocks was required; additionally, for short-read alignments containing splice junctions, exact-match junctions supported by long reads on the same strand were required. Parameter values used for mask construction and candidate selection were calibrated on a reference set of well-characterized factors (SRSF1, SRSF2 and hnRNPM) and then fixed for all RBPs analyzed in this study (**Extended Data Fig.** 2a–e). These parameters included the long-read length upper bound for mask construction, the minimum overlap length between short-read alignments and long-read exon blocks, and the minimum long-read support required to define exon/junction evidence. Long-read alignments longer than 500 nt were excluded from mask construction to avoid overly broad exon masks that could reduce positional resolution for short-read attribution. A minimum short-read/long-read exon-block overlap of 40 nt was required to ensure substantial overlap between candidate short-read placements and long-read-supported transcript structure. A minimum long-read support threshold of one was used to retain sensitivity in regions with sparse long-read CLIP coverage. Using the calibrated thresholds, the candidate placement with the maximum overlap between the short read and long-read exon blocks was selected. Ties were resolved by the highest mean long-read per-base depth across the overlapped bases, and remaining ties were resolved deterministically to ensure reproducibility. Multimapping reads for which none of the candidate placements satisfied these criteria were discarded. This procedure was implemented in custom scripts and produced filtered short-read BAMs.

### CITS quantification and peak calling

Filtered BAMs were UMI-deduplicated with UMIcollapse and processed with the Skipper pipeline^25^ under default settings. ENCODE eCLIP-derived blacklist filters were not directly applied to SCALE-CLIP datasets because SCALE-CLIP used an anti-FLAG antibody rather than heterogeneous RBP-specific antibodies.

### Analysis of ENCODE eCLIP

eCLIP datasets used in this study are listed in Supplementary Data 7. They were processed using the same genome build, transcript annotation, analysis pipeline (Skipper)^25^ and FDR definition as used for SCALE-CLIP. For ENCODE datasets, blacklist filtering implemented in Skipper was enabled. To compare reproducibility, pairwise Jaccard similarity was computed on the intersection of tested transcript windows shared by the replicate pair. For three-replicate SCALE-CLIP datasets, Jaccard was computed for all three replicate pairs and summarized by the median.

### 5-Ethynyluridine (5-EU) pulse-chase-coupled CLIP-seq

#### Experimental methods

For each replicate, cells were prepared at a scale of 20 million cells per condition. For pulse labeling, cells were incubated with 2 mM 5-EU (TCI, E1560) for 10 min, a short pulse chosen to minimize potential cytotoxicity. After centrifugation (500 × g, 3 min), the medium was replaced with RPMI 1640 containing 4 mM uridine for a chase of 10- or 30-min. Pulse-chase labeling conditions were adapted from the prior study^54^ with modifications (5-EU in place of 4sU and increased nucleoside concentrations) to improve incorporation and recovery of labeled RNA for downstream CLIP. Cells were washed with ice-cold PBS and UV-crosslinked on ice at 400 mJ/cm² (254 nm). Crosslinked cells were lysed, fragmented, immunoprecipitated, and resolved by SDS-PAGE; RNA fragments were recovered from nitrocellulose as described in the “Short-read seCLIP” procedure of the long-read-guided enhanced CLIP-seq section^49^. Recovered RNA fragments were then subjected to Cu(I)-catalyzed azide-alkyne cycloaddition (CuAAC) click chemistry to selectively capture 5-EU-containing RNAs, followed by streptavidin enrichment and reverse transcription using the Click-iT Nascent RNA Capture Kit (Invitrogen, C10365) according to the manufacturer’s instructions. Resulting cDNAs were converted to sequencing libraries following standard seCLIP protocols and sequenced on an Illumina NextSeq (single-end, 75 bp), yielding ≥ 20 million reads per replicate. Each experiment included two biological replicates with size-matched input controls.

#### Data processing

Primary processing was performed with the Skipper pipeline^25^ with default parameters. Reads were aligned to reference human genome (GRCh38, no-alt-analysis set) with GENCODE v44 annotation using STAR (v2.7.10b) ^52^. Skipper then partitioned each transcript into non-overlapping windows of up to 100 nt (shared across samples) and counted CITS for IP and size-matched input libraries within each window. For temporal grouping, windows called as reproducibly enriched at least once in each RBP dataset were retained. Reproducible enrichment calls at each time point were binarized, with 1 denoting reproducible enrichment and 0 denoting no reproducible enrichment call. Windows were classified by their binary pattern across (SRSF1 10 min, SRSF1 30 min, SRSF6 10 min, SRSF6 30 min): reciprocal switch, (1,0,0,1) or (0,1,1,0); single-factor sustain, (1,1,0,1), (1,1,1,0), (1,0,1,1), or (0,1,1,1); coordinated change, (1,0,1,0) or (0,1,0,1); and dual sustain, (1,1,1,1).

### KD-CLIP-seq

#### Experimental methods

siRNAs were transfected with Lipofectamine RNAiMAX (Invitrogen) according to the manufacturer’s protocol. For all transfections, siRNAs were used at a final concentration of 40 nM (Silencer Select) or 20 nM (DsiRNA and siGENOME). Knockdown efficiency was verified by RT-qPCR, with ≥ 60% reduction of target transcript relative to the control sample considered acceptable. Knockdowns were performed in biological duplicate, and siRNA sequences are listed in Supplementary Data 6. After 48 h, siRNA-transfected cells were UV crosslinked on ice at 400mJ/cm^2^ with 254 nm radiation. A portion of the cells was collected separately for RNA extraction using TRIzol Reagent (Invitrogen) and Direct-zol RNA Miniprep kit (Zymo Research, R2052), and knockdown efficiency was verified by RT-qPCR. qPCR Primer sequences are listed in Supplementary Data 6. CLIP libraries were prepared following the “Short-read seCLIP” procedure, adapted from the original seCLIP protocol^49^.

#### Differential binding analysis

Primary processing of seCLIP data was performed with the Skipper pipeline^25^ using default settings. Count tables (IP and input) from control and knockdown conditions were merged by window to form a unified matrix comprising windows detected in either condition. Windows absent from a given sample were assigned zero counts. Differential binding was assessed in edgeR (v4.0.16)^55^. Low-count windows were filtered using filterByExpr (and a minimal CPM threshold to reduce low-count artifacts, applied to both IP and input libraries), and library sizes were normalized by the trimmed mean of M values (TMM). A negative binomial generalized linear model was fitted with design ∼ condition * sample_type, where the interaction term tests whether IP-over-input enrichment differs between control and knockdown. Significance was evaluated using quasi-likelihood F tests (glmQLFit with robust=TRUE; glmQLFTest). Windows with FDR < 0.05 and |log2FC| > 1 were considered differentially bound, where log2FC represents the change in IP-over-input enrichment in knockdown relative to control. Sample relationships were visualized by MDS on logCPM values (edgeR plotMDS).

### Chromosome-associated RNA (caRNA) and nuclear RNA extraction

20 million cells were fractionated using NE-PER Nuclear and Cytoplasmic Extraction Reagents (Thermo Scientific, 78833) following manufacturer’s instructions. For total nuclear RNA extraction, the nuclear pellet was resuspended in TRIzol reagent, homogenized by gentle pipetting, and RNA was purified using Direct-zol RNA Miniprep kit (Zymo Research, R2052). Chromosome-associated RNA (caRNA) was isolated based on a previous study^56^ with minor modifications. Briefly, NE-PER-derived nuclear fraction was centrifuged at 4 °C at 15,000 × g for 2 min to separate the soluble nuclear (supernatant) and chromosome-associated (pellet) fractions. The supernatant containing the soluble nuclear fraction was discarded. The chromosome-associated pellet was washed twice with cold PBS containing 1 mM EDTA, resuspended in TRIzol reagent, homogenized by gentle pipetting, and RNA was purified using Direct-zol RNA Miniprep kit (Zymo Research, R2052).

### Direct RNA sequencing (ONT) for transcriptome-wide RNA modification profiling

#### Experimental methods

Nuclear RNA or chromosome-associated RNA extracted following ‘Chromosome-associated RNA (caRNA) and nuclear RNA extraction’ section was depleted of ribosomal RNA using RiboMinus Eukaryote Kit v2 (Invitrogen, A15020). rRNA depletion efficiency was assessed by Tapestation. To enable ONT direct RNA sequencing of nascent RNAs lacking poly(A) tails, rRNA-depleted RNA was polyadenylated using poly(A) polymerase (Takara, 2180A) at 37°C for 15 min, followed by purification with RNA Clean & Concentrator-5. Sequence libraries were prepared with the Direct RNA Sequencing Kit (SQK-RNA004, Oxford Nanopore Technologies) according to the manufacturer’s protocol and sequenced on a PromethION instrument using Flow Cell FLO-PRO004RA (ONT).

#### Data processing

The raw Nanopore RNA-seq data (POD5) were basecalled using Dorado (v1.0.1) with the model rna004_130bps_hac@v5.2.0, with modified basecalling enabled (m⁵C, inosine/m⁶A, and pseudouridine (Ψ)). Basecalled reads were aligned to human reference genome (GRCh38) using minimap2 (v2.28)^53^ with the parameters -x splice -k14. Base modification calls were summarized and aggregated into site-level metrics using modkit (v0.5.0). For downstream analysis, base positions in the resulting BED files were filtered to retain only those with (i) valid canonical base coverage ≥ 20 and (ii) canonical-to-total basecalling ratio > 0.7. To define a higher-confidence m^6^A site set for downstream analyses, the Dorado m^6^A calls was orthogonally benchmarked against an independent K562 caRNA MeRIP-seq dataset (GSE205709)^56^. Sites with an estimated m^6^A modification fraction of ≥ 30% showed substantially greater overlap with reproducible MeRIP-seq peaks (Extended Data Fig. 9a–c), and this threshold was used for subsequent analyses unless otherwise stated.

### KD-RNA-seq

#### Experimental methods

siRNAs targeting each RBP were obtained from IDT (DsiRNA), Invitrogen (Silencer Select) or Dharmacon (siGENOME). siRNAs were transfected into parental K562 cells with Lipofectamine RNAiMAX (Invitrogen) according to the manufacturer’s protocol. For all transfections, siRNAs were used at a final concentration of 40 nM (Silencer Select) or 20 nM (DsiRNA and siGENOME). Knockdowns were performed in biological triplicate. To minimize batch effects, a matched non-targeting control siRNA was included for each transfection batch and experimental time point, with separate controls used for each siRNA product type. After 48 h, total RNA was extracted using TRIzol Reagent (Invitrogen) and Direct-zol RNA Miniprep kit (Zymo Research, R2052). Knockdown efficiency was verified by RT-qPCR, with ≥ 60% reduction of target transcript relative to the control sample considered acceptable. siRNA sequences and primer sequences are listed in Supplementary Data 6. For short-read RNA sequencing, libraries were prepared using the DNBSEQ Eukaryotic Strand-specific Transcriptome Resequencing protocol (DNBSEQ stranded mRNA library) and sequenced on the DNBSEQ platform to generate paired-end 100bp reads, yielding ≥ 20 million reads per replicate. One of three replicates was also used for long-read RNA-seq. For long-read RNA-seq, cDNA libraries were prepared with the PCR-cDNA Sequencing Kit (SQK-PCS114, Oxford Nanopore Technologies) following the manufacturer’s protocol and sequenced on a PromethION instrument using Flow Cell R10 (FLO-PRO114M, ONT).

#### Primary data processing

For short-read RNA-seq, reads were aligned to the human reference genome GRCh38 (no-alt analysis set; GCA_000001405.15) using the GENCODE v44 annotation with STAR (v2.7.10b)^52^ in two-pass mode and default parameters. For long-read RNA-seq, raw pod5 files were base-called with Dorado version 1.0.1 (ONT) in high-accuracy mode. Reads were subsequently filtered (Q ≥ 9, read length ≥ 100 bp) and oriented using Pychopper (v2.7.10), and only full-length oriented reads were aligned to GRCh38 using minimap2 (v2.28; parameters: -ax splice -uf -k14 - secondary=no)^53^, and alignments with MAPQ ≥ 20 were retained for downstream analysis.

#### Hybrid transcript assembly

A hybrid transcript annotation was generated using StringTie (v3.0.0)^57^ by merging the GENCODE v44 annotation with StringTie-assembled transcripts derived from paired long-read and short-read RNA-seq data (parameters: --mix -p 4 -f 0.001 -c 0.5 -t). Hybrid assemblies independently generated for each SF knockdown and control knockdown conditions were subsequently merged with StringTie (-p 4 -i -G gencode.v44.annotation.gtf -F 0.1 -T 0.1 -f 0.001). The resulting merged annotation was quality-assessed and filtered with SQANTI3 (v5.5.1)^58^. A genome index was then re-generated based on SQANTI3-filtered hybrid transcript annotation, and short-read RNA-seq reads were re-aligned using STAR. The resulting alignments were used for downstream analyses.

#### Gene expression quantification

Transcript abundance was quantified using StringTie (v3.0.0). Differential gene expression analysis was performed with edgeR (v4.0.16)^55^ after TMM normalization. Genes with FDR < 0.05 and absolute log_2_(fold-change) > 1 were considered differentially expressed.

#### Splicing quantification

Differential splicing events were analyzed using rMATS-turbo (v4.1.2)^59^. Unless otherwise stated, rMATS results based on the hybrid transcript annotation were used for downstream analyses. rMATS-turbo identified five major classes of alternative splicing (AS) events: skipped exon (SE), mutually exclusive exons (MXE), alternative 3’ splice site (A3SS), alternative 5’ splice site (A5SS) and retained intron (RI). Events with absolute inclusion level difference > 0.05 and FDR < 0.05 were considered significant differential splicing events unless otherwise specified.

### Immunofluorescence

3×FLAG-tagged K562 cells were seeded onto Poly-D-Lysine coated German glass coverslips (18 mm round, #1 thickness; NEU, H-18-PDL) in 6-well or 12-well plates. After cell attachment, cells were fixed with 4% paraformaldehyde in PBS for 15 min, permeabilized in 0.5% Triton X-100 in PBS for 30 min and blocked with 3% BSA in PBS (filtered through 0.22 µm PVDF membrane) for 1 h, all conducted at room temperature. Primary antibodies (mouse anti-SC35, ab11826, 1:500, Abcam; rabbit anti-DDDDK tag, ab205606, 1:300, Abcam) diluted in 3% BSA were applied to the cells and incubated overnight at 4 °C. After three washes with PBST for 10 min each, cells were incubated for 1 h at room temperature with secondary antibodies (Goat Anti-Rabbit IgG H&L Alexa Fluor 488, ab150077, 1:1000, Abcam; Goat Anti-Mouse IgG H&L Alexa Fluor 568, ab175473, 1:1000, Abcam). After three PBST washes, nuclei were counterstained with Hoechst 33342 (2 µg/mL in PBS) for 30 min at room temperature and mounted with ProLong Gold Antifade Mountant (Invitrogen, P36930). Images were acquired on a FLUOVIEW FV3000 Confocal Laser Scanning Microscope (Olympus / Evident) equipped with a 60× oil immersion objective. Raw images were processed using Fiji/ImageJ (1.54p). All images shown were subjected to identical adjustments of brightness/contrast within an experiment.

### RNA oligonucleotide binding assay

5’-Biotinylated RNA oligonucleotides containing a common SRSF-binding motif (GAAGA) and a 3’-dTdT tail were synthesized by IDT. The oligonucleotide sequence is provided in Supplementary Data 6. RNA oligos were incubated overnight at 4 °C with RNA-fragmented FLAG-tagged K562 cell lysates to allow SRSF binding. The lysis buffer was identical to that used in CLIP experiments (50 mM Tris-HCl pH 7.4, 100 mM NaCl, 1% (vol/vol) Igepal CA-630, 0.1% (vol/vol) SDS and 0.5% (wt/vol) sodium deoxycholate) supplemented with protease and phosphatase inhibitors (Millipore, 539134; Sigma Aldrich, P0044). The reaction mixture was UV crosslinked on ice at 400 mJ/cm^2^ with 254 nm radiation, and immunoprecipitated with anti-FLAG M2 Magnetic Beads (Millipore, M8823) or mouse IgG1 (isotype control)-Magnetic Beads (MBL, M075-11) for 2-3 hours at 4 °C. After extensive high- and low-salt washes, immunoprecipitated FLAG-tagged RBP and bound RNA oligos were eluted with SDS sample buffer and separated by SDS-PAGE. FLAG-tagged SFs were visualized by Western blot with DYKDDDDK tag Polyclonal antibody (Proteintech, 20543-1-AP) and biotinylated RNA oligos were visualized by avidin HRP with Chemiluminescent Nucleic Acid Detection Module Kit (Thermo Scientific, 89880). High-salt wash buffer contained 50 mM Tris-HCl pH 7.4, 1 M NaCl, 1% (vol/vol) Igepal CA-630, 1 mM EDTA, 0.1% (vol/vol) SDS and 0.5% (wt/vol) sodium deoxycholate. Low-salt wash buffer contained 20 mM Tris-HCl pH 7.4, 10 mM MgCl_2_, 0.2% (vol/vol) Tween-20 and 5 mM NaCl.

### Immunoprecipitation-mass spectrometry (IP-MS)

For proteomic analysis of SRSF-associated complexes, SRSF1–11 3×FLAG-tagged and parental K562 cells were lysed in 1% digitonin in 1× TBS buffer supplemented with protease and phosphatase inhibitors. After clarification by centrifugation, the lysates were incubated for 1 h at 4 °C with Dynabeads Protein G (Thermo Fisher Scientific, DB10004) coupled to anti-FLAG M2 antibody (Sigma-Aldrich, F3165). The beads were then washed extensively with 1× TBS containing 0.1% Triton X-100, and bound proteins were eluted with FLAG peptide. Eluted proteins were precipitated with trichloroacetic acid (TCA) and resuspended in 7 M guanidine hydrochloride with 0.1 M ammonium bicarbonate (pH 8.8) containing 0.05% decyl glucoside. Samples were reduced with TCEP and alkylated with iodoacetamide, then diluted to a final guanidine hydrochloride concentration of 1.4 M and digested with Lys-C at 37 °C for 2 h. The samples were subsequently further diluted to a final guanidine hydrochloride concentration of 0.7 M and digested with trypsin overnight at 37 °C. The resulting peptides were acidified with trifluoroacetic acid and desalted using EvoTips prior to LC-MS/MS analysis. Peptides were analyzed on an Orbitrap Astral mass spectrometer coupled to an Evosep One system. Each sample was analyzed in two technical replicates. MS data were processed using DIA-NN v1.8 against the UniProt human reference proteome (UP000005640), with peptide- and protein-level false discovery rates controlled at 1%. To quantify SRSF-associated protein enrichment, protein enrichment was calculated for each bait replicate as log_2_((bait signal + 1)/(mean parental-control signal + 1)). Replicate-level enrichment scores were averaged for each bait. Proteins were annotated into spliceosome-related functional classes using the curated lists of spliceosomal complex components provided in Supplementary Table 1 of the reference paper ^60^. For class-level visualization, each bait–class pair was summarized by the median protein-level mean log2 enrichment of quantified class members.

### Oligo pull-down assay

5′-Biotinylated RNA oligonucleotides were synthesized by IDT (sequences in Supplementary Data 6). For the m^6^A-dependent binding assay, 20–30 nt RNA oligonucleotides were designed to span a called m^6^A site and an adjacent SRSF motif within each selected SCALE-CLIP peak. Matched A and m^6^A oligonucleotide pairs differing only at the modified nucleotide were synthesized for each locus (Supplementary Data 6). All loci also fall within reproducible K562 caRNA MeRIP-seq intervals from an independent dataset (GSE205709)^56^. RNA oligos (0.8 nmol) were incubated with NeutrAvidin Agarose beads (Pierce/Thermo Scientific, 29200; 100 µL slurry) in 1×TBS for 2 h at 4 °C with rotation to immobilize biotinylated RNA. Beads were washed three times with 1×TBS. K562 cells endogenously expressing 3×FLAG-tagged RBPs were lysed in lysis buffer containing 1% digitonin in 1×TBS supplemented with Protease Inhibitor Cocktail III (Millipore, 539134), Phosphatase Inhibitor Cocktail 3 (Sigma-Aldrich, P0044) and RNase Inhibitor, Murine (NEB, M0314) followed by sonication and centrifugation at 15,000×g for 10 min at 4 °C. Clarified lysates (20 million cells/ml) were incubated with RNA-coated beads in a total volume of 200 µL overnight at 4 °C with rotation. Where indicated, RNA-coated beads were pre-incubated with ASOs (phosphorothioate backbone and 2′-O-methyl modification; synthesized by IDT; sequences in Supplementary Data 6) at a final concentration of a 1, 2 or 5 µM for 10 min at 4 °C prior to addition of cell lysates. A non-targeting control ASO was included in parallel. Beads were then washed three times with 1×TBS. Bound proteins were eluted by boiling in SDS sample buffer at 95 °C for 5 min and analyzed by SDS-PAGE and immunoblotting using anti-FLAG M2 antibody (Sigma-Aldrich, F1804).

### SCALE-CLIP-guided Antisense oligonucleotide design and transfection

Antisense oligonucleotides (ASOs; phosphorothioate backbone and 2′-O-methyl modification) were synthesized by IDT. ASO sequences are listed in Supplementary Data 6. To block SRSF binding and modulate splicing, ASOs were designed within SCALE-CLIP-defined SRSF binding peaks overlapping splicing-altered regions and positioned to cover enriched SRSF binding motifs within each peak. For each targeted peak, one or two independent ASOs were designed to target distinct motif instances and/or adjacent positions within the peak. HeLa or K562 cells were transfected with ASO using Lipofectamine 3000 (Invitrogen) according to the manufacturer’s instructions. ASOs were tested at final concentrations of 0.05, 0.1, and 0.3 µM. A non-targeting control ASO was included in each experiment at final concentrations of 0.3 µM. Cells were harvested 24 h post-transfection, and total RNA was extracted for RT-PCR analysis followed by gel electrophoresis. PCR products were then quantified by TapeStation where indicated. For ASO transfection experiments targeting the EZH2 NMD-associated exon, cells were treated with cycloheximide (0.1 mg/ml) for 5 h before harvest to suppress nonsense-mediated decay.

### RT-PCR and qPCR

Total RNA was extracted using TRIzol reagent (Invitrogen) and Direct-zol RNA Miniprep kit (Zymo Research, R2052) according to the manufacturer’s instructions. Typically, 0.2-1 µg of total RNA was reverse transcribed with ReverTra Ace qPCR RT Master Mix (Takara) to generate cDNA. For semi-quantitative RT-PCR, cDNA corresponding to 10-50 ng RNA was amplified with ExTaq DNA polymerase (Takara) using target-specific primer pairs, and products were resolved by agarose-gel electrophoresis and visualized by SAFELOOK green nucleic acid stain (FUJIFILM). Primer sequences used for RT-PCR are listed in Supplementary Data 6. For RT-qPCR, reactions were performed in technical triplicates using SYBR Green Realtime PCR Master Mix (TOYOBO, QPK-201) on a CFX Duet Real-Time PCR System (Bio-Rad). Relative mRNA levels were calculated by the 2^-ΔΔCt^ method using GAPDH as the internal control. qPCR primer sequences are listed in Supplementary Data 6.

### Western blotting

Cells were lysed and denatured with Tris-SDS sample buffer with 2-mercaptoethanol (Nacalai, 30566-22) and separated by SDS-PAGE and transferred to PVDF or nitrocellulose membranes. Primary antibodies used for Western blotting are as follows: anti-FLAG M2 mouse monoclonal antibody (Sigma-Aldrich, F1804) at 1:1000; DYKDDDDK tag rabbit polyclonal antibody (Proteintech, 20543-1-AP) at 1:2000; and GAPDH (14C10) rabbit monoclonal antibody (CST, 2128L) at 1:1000.

### Motif analysis

Motif enrichment analyses were performed on SCALE-CLIP fine-mapped binding peaks using HOMER findMotifsGenome.pl (v4.11.1) with default settings from Skipper pipeline. For additional analyses outside the Skipper, the same HOMER settings were used (-size given -rna -nofacts -S 20-len 5,6,7,8,9 -nlen 1), while background regions were generated automatically by HOMER. Motif occurrences within peaks were identified using annotatePeaks.pl with a predefined motif PWM.

### Repeat element annotation and enrichment analysis of CLIP peaks

Repeat annotations were obtained from the UCSC Genome Browser RepeatMasker track (rmsk, hg38/GRCh38). SCALE-CLIP fine-mapped binding peaks were intersected with RepeatMasker annotations using bedtools intersect (v2.31.1) with -wao to obtain overlap lengths. When multiple repeats overlapped a peak, a single repeat was assigned per peak by prioritizing larger overlap length. Peaks with no overlap were labeled No_repeat. Opposite-strand overlaps were retained for LINE/SINE/LTR and labeled as antisense (_AS).

### Meta-exon profiling

Meta-exon profiles were generated for cassette exon (SE) events using the rbp-maps pipeline^28^. Cassette exon annotations were obtained from rMATS-turbo SE output files (SE.MATS.JCEC) generated from all control and target RBP knockdown samples with three biological replicates per condition. To remove redundant event entries, each rMATS SE file was processed with subset_jxc (rbp-maps container; docker://brianyee/rbp-maps:a5c9bd9) using -e se. Overlapping alternative splicing event regions were collapsed, and a single representative event was retained per merged region by selecting the event with the highest average inclusion junction count, calculated across all replicates and conditions. Meta-exon profiles were computed by aggregating peak occurrences around SE exon-intron boundaries using rbp-maps (plot_map) with an exon flank of 50 nt and an intron flank of 500 nt (--exon_offset 50 --intron_offset 500). The per-position histogram output (hist.txt) produced by rbp-maps was used for downstream visualization and cross-RBP comparisons.

### Integrative analysis of CLIP peaks and KD-RNA-seq

To associate splicing regulation with RBP binding patterns, SE events identified from KD-RNA-seq were stratified according to knockdown-induced splicing changes. Significant differential SE events were defined as FDR < 0.05 and |ΔPSI| > 5%. Binding profiles around differentially spliced regions were then extracted as described above using rbp-maps separately for exons included upon knockdown (ΔPSI > 5%) and skipped upon knockdown (ΔPSI < −5%). For each RBP, SE events with at least one fine-mapped SCALE-CLIP peak for the corresponding RBP within the meta-exon profiling window were used to visualize differences in positional binding patterns between exons included and skipped upon RBP knockdown.

### SpliceAI scoring of splice sites

Splice site probability scores were estimated with SpliceAI^61^. SpliceAI (v1.3.1) was run in custom sequence mode to score splice sites within regions showing knockdown-responsive splicing changes. For each splicing junction, the strand-aware genomic sequence spanning the annotated site with ±5,000 bp context (total 10 kb) was extracted. The five pretrained SpliceAI-10k models were loaded and the ensemble mean prediction across models was computed. From the network output, splice site probabilities of the acceptor and donor site of alternative exon and flanking exons were extracted. All steps were implemented in Python using the SpliceAI API (keras, spliceai.utils.one_hot_encode) and run with default network weights.

### Multivariable modelling of baseline exon inclusion and cognate SRSF perturbation sensitivity

To model steady-state exon inclusion rate, linear regression models were fitted to baseline PSI values using cassette exons with suboptimal splice sites. Alternative-exon SRSF multiplicity, upstream and downstream flanking-exon SRSF multiplicity, SpliceAI-10k splice-site scores for donor and acceptor site of alternative exon, and alternative exon length were included as predictors. Baseline PSI and continuous variables were z-scored before model fitting. To model perturbation sensitivity, logistic regression models were fitted across exon–SRSF pairs. For exon-centered analyses, cassette exons lacking detectable SRSF binding on the flanking exons were selected, and cognate SRSF binding on the alternative exon was classified as none, target only, or target plus others (at least one additional SRSF family member). The response variable was defined as significant knockdown-induced skipping. For flanking analyses, cassette exons lacking detectable SRSF binding on the alternative exon were selected, and cognate SRSF binding on the flanking exons was classified as none, target only on either flanking exon, or target plus others on either of the flanking exons. The response variable was defined as significant knockdown-induced inclusion. In both perturbation models, splice-site strength, baseline PSI, and alternative exon length were included as covariates.

### Analysis of RNA modification correlation with SCALE-CLIP peaks

RNA modification sites for m^6^A, inosine, pseudouridine (ψ), and m^5^C were called from ONT direct RNA-seq as described above. Sites with coverage ≥ 20 and canonical-to-total basecalling ratio > 0.7 were retained and intersected with strand-matched SCALE-CLIP reproducible enriched windows. For each RBP and modification type, sites were stratified into high (≥ 30%) and low (< 30%) modification groups, and enrichment of SCALE-CLIP overlap in the high group was assessed using Fisher’s exact test with BH-FDR correction across RBPs. Sites were also binned by percent modification in 10% increments, and the fraction overlapping SCALE-CLIP windows was calculated per bin. Binned overlap-rate profiles were column-normalized across modification bins for each RBP before heatmap plotting.

For motif-centered analyses, HOMER-defined top enriched motifs in SCALE-CLIP fine-mapped peaks were used as anchor motifs. Motif occurrences were identified using annotatePeaks.pl, and high-confidence m^6^A sites were defined as sites with modification frequency ≥ 30%, read depth ≥ 20, and canonical A-call fraction ≥ 0.7. Motif-centered metaprofiles were generated by calculating the fraction of motif occurrences containing a high-confidence m^6^A site at each strand-aware offset from −30 to +30 nt relative to the motif center. For even-length motifs, position 0 was defined as the downstream of the two central nucleotides. Profiles were normalized to a maximum value of 1. For motif-internal analyses, high-confidence m^6^A sites within motif bodies were used to calculate the relative positional distribution of m^6^A sites, grouped by motif length.

### Cross-cell-type analysis of m^6^A–splicing coupling

Matched m^6^A levels and exon inclusion rates were analyzed across K562, KG-1a, Nalm-6, HEL, HeLa, A549 and HepG2 cells. Nuclear RNA m^6^A levels were quantified using Dorado and modkit as described above, and splicing was quantified from total RNA-seq using rMATS-turbo. For each RBP, eligible site-event pairs were defined using K562-derived SCALE-CLIP peaks as anchors and consisted of an adenosine site that passed the m^6^A modification-calling filters, overlapped a strand-matched SCALE-CLIP peak, and was linked to the corresponding cassette exon. The same genomic m^6^A site and cassette exon were queried across cell types, with cell types lacking either a valid m^6^A estimate or PSI measurements were treated as missing. For each site-event pair, Spearman’s ρ was calculated across cell types using percent m^6^A and PSI, retaining only pairs measured in at least four cell types. Correlation coefficients were summarized for each RBP. For cell-type-stratified analysis, each RBP-linked site-event pair was stratified within each cell type by m^6^A level (< 30% or ≥ 30%). PSI values were logit-transformed after bounding to [0.01, 0.99], and low- and high-m^6^A groups were compared using a two-sided Mann-Whitney U test.

### Statistics and reproducibility

No statistical method was used to predetermine sample size. Experiments were not randomized, and investigators were not blinded to group allocation during data acquisition or analysis unless otherwise stated. Number of biological or technical replicates are described in the corresponding Methods sections. Unless otherwise stated, statistical tests were two-sided. Statistical methods for individual analyses are described in the corresponding Methods sections and figure legends. Multiple testing correction was performed using the Benjamini–Hochberg procedure unless otherwise stated. Exact P values, FDR values, test statistics, effect sizes and n values are provided in the figure, figure legends, Source Data or Supplementary Data where applicable. Error bars and box-plot elements are defined in the corresponding figure legends. Representative images, immunoblots and gels are shown from experiments repeated independently as stated in the corresponding figure legends.

## Reporting summary

Further information on research design is available in the Nature Portfolio Reporting Summary linked to this article.

## Data availability

Sequencing data generated in this study have been deposited in the Gene Expression Omnibus (GEO), with raw sequencing data available through the Sequence Read Archive (SRA). The mass spectrometry data have been deposited in the Japan ProteOme STandard Repository (jPOST) via the ProteomeXchange Consortium. All other RNA-seq datasets used in this study are listed in Supplementary Data 7.

## Code availability

Custom scripts used for long-read-guided read attribution (LRA) in SCALE-CLIP are available at https://github.com/minayoshida/scaleclip-lra. An archived release with a DOI will be provided upon publication.

## Acknowledgements

We thank all members of the Yoshimi lab for helpful discussions and technical support. The supercomputing resource, SHIROKANE, provided by Human Genome Center was used for large data processing (the University of Tokyo). M.Y. was supported by JSPS Research Fellowship for Young Scientists. This work was supported by JSPS KAKENHI Grants-in-Aid for Scientific Research (B) (JP23K27379) to M.A.; grants from the Takeda Science Foundation to M.A. and A.Y.; MEXT KAKENHI for Transformative Research Areas (A) (JP23H04414) to R.S.K.; Cabinet Office BRIDGE (programs for bridging the gap between R&D and the ideal society (Society 5.0) and generating economic and social value) to R.H.; JST CREST (JPMJCR20E6 and JPMJCR23B3) and AMED-CREST (JP23gm1610010) to S.A.; BINDS from AMED; JSPS KAKENHI Grants-in-Aid for Scientific Research (A) (JP21H04828 and JP26H02447) to A.Y.; Research on Development of New Drugs from AMED (JP26ak0101275 and JP26ak0101263) to A.Y.; Practical Research for Innovative Cancer Control from AMED (JP26ck0106946 and JP26ck0106906) to A.Y.; AMED-PRIME (JP26gm7010010) to A.Y.; JST FOREST (JPMJFR2118) to A.Y.; the G-7 Scholarship Foundation and the Canon Foundation to A.Y.

## Author Contributions

M.Y., M.A. and A.Y. designed the study. M.Y, K.N., R.M., M.H., N.S., M.S., S.K., and R.H. performed experiments. M.Y. performed the computational analyses. M.Y., M.A. and A.Y. interpreted the data. S.A. performed mass-spectrometry data acquisition and analysis. R.S.K. assisted with microscopy analysis. H.U., G.N., and H.A. assisted with long-read RNA-seq analysis. M.A., H.M., and A.I. provided guidance on experiments and computational analyses. A.Y. supervised the study. M.Y., M.A., and A.Y. wrote the first draft of the manuscript. M.Y., M.A. and A.Y. reviewed and edited the manuscript. All authors reviewed the manuscript and approved the final version.

## Extended Data Figures and Legends

**Extended Data Fig. 1.**
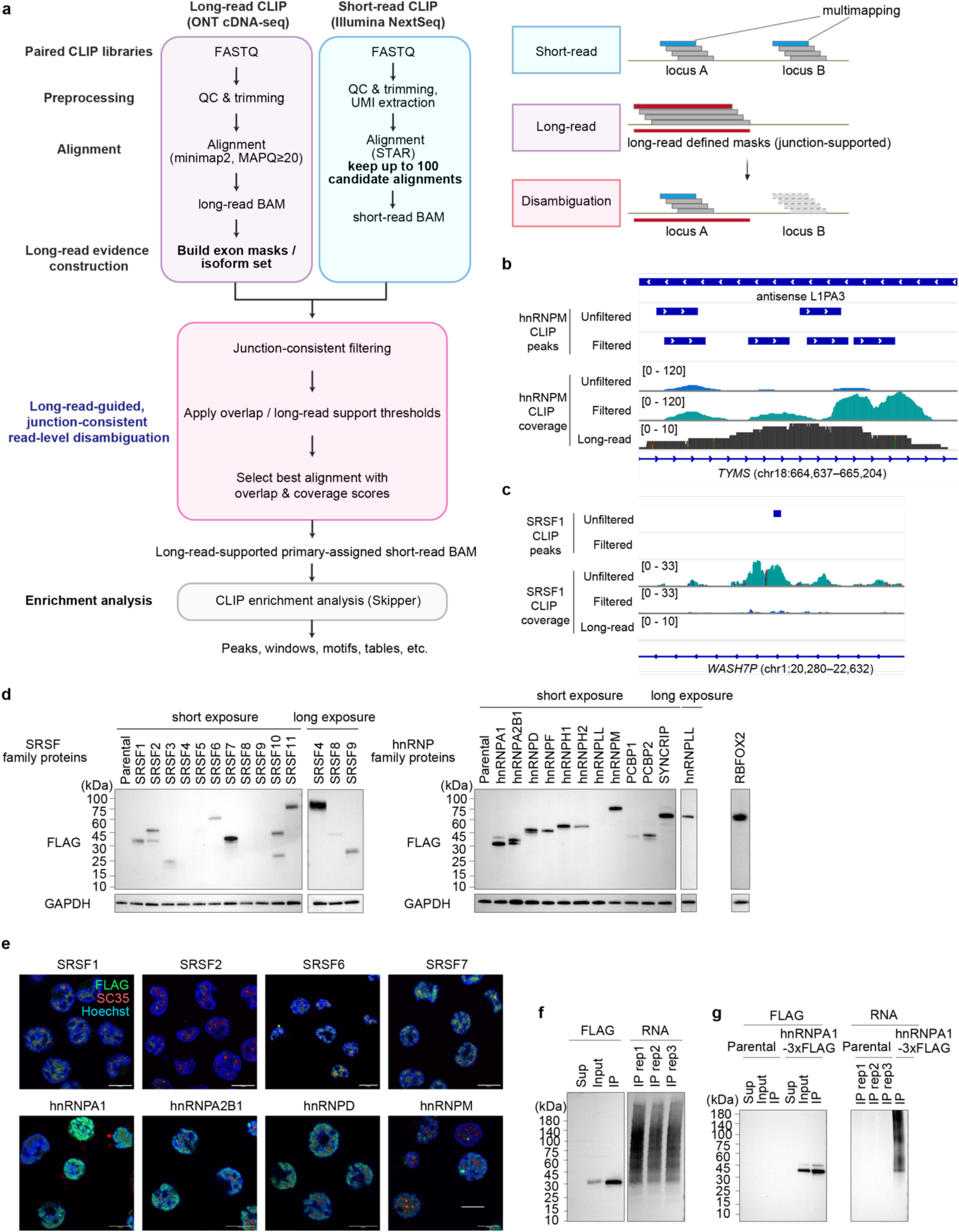
Computational pipeline for SCALE-CLIP and quality control for endogenously tagged cells. **a,** Computational pipeline for long-read-guided read attribution (LRA). Short-read CLIP reads were aligned allowing up to 100 multi-mapped positions, while long reads were uniquely mapped. Short-read alignments were filtered based on consistency with long-read-derived splice junctions and overlap with long-read-supported exonic regions, and the alignment with the largest overlap and highest long-read coverage when tied was selected. Filtered short-read BAM files were then analyzed with the Skipper pipeline. **b, c,** Representative genome browser views showing the effect of LRA on short-read CLIP coverage and peak detection. **b,** Within TYMS overlapping an antisense L1PA3 element, LRA redistributed short-read hnRNPM CLIP coverage to long-read-supported regions, resulting in detection of an LRA-specific peak. **c,** Within WASH7P, LRA reduced short-read SRSF1 CLIP coverage in regions lacking long-read support, resulting loss of the peak detected without LRA. Coverage tracks with and without LRA are shown on the same y-axis scale. **d,** Immunoblot of 3×FLAG-tagged SRSFs (left) and hnRNPs (right) used in SCALE-CLIP experiments. For SRSF4, SRSF8, SRSF9 and hnRNPLL, longer-exposure images are shown to better visualize low-expression FLAG signal. **e,** Representative immunofluorescence image of selected high-signal 3×FLAG knock-in cell lines stained for FLAG (green), SC35 (red), and Hoechst (blue). Scale bar, 10 µm. **f,** Representative IP-Western blot (left) and biotin-labelled RNA fragments recovered from SCALE-CLIP (right) for SRSF1-3×FLAG cells. **g,** Representative IP-Western blot (left) and recovered RNA (right) from parental K562 and hnRNPA1-3×FLAG cells, showing minimal background recovery in the parental control.

**Extended Data Fig. 2.**
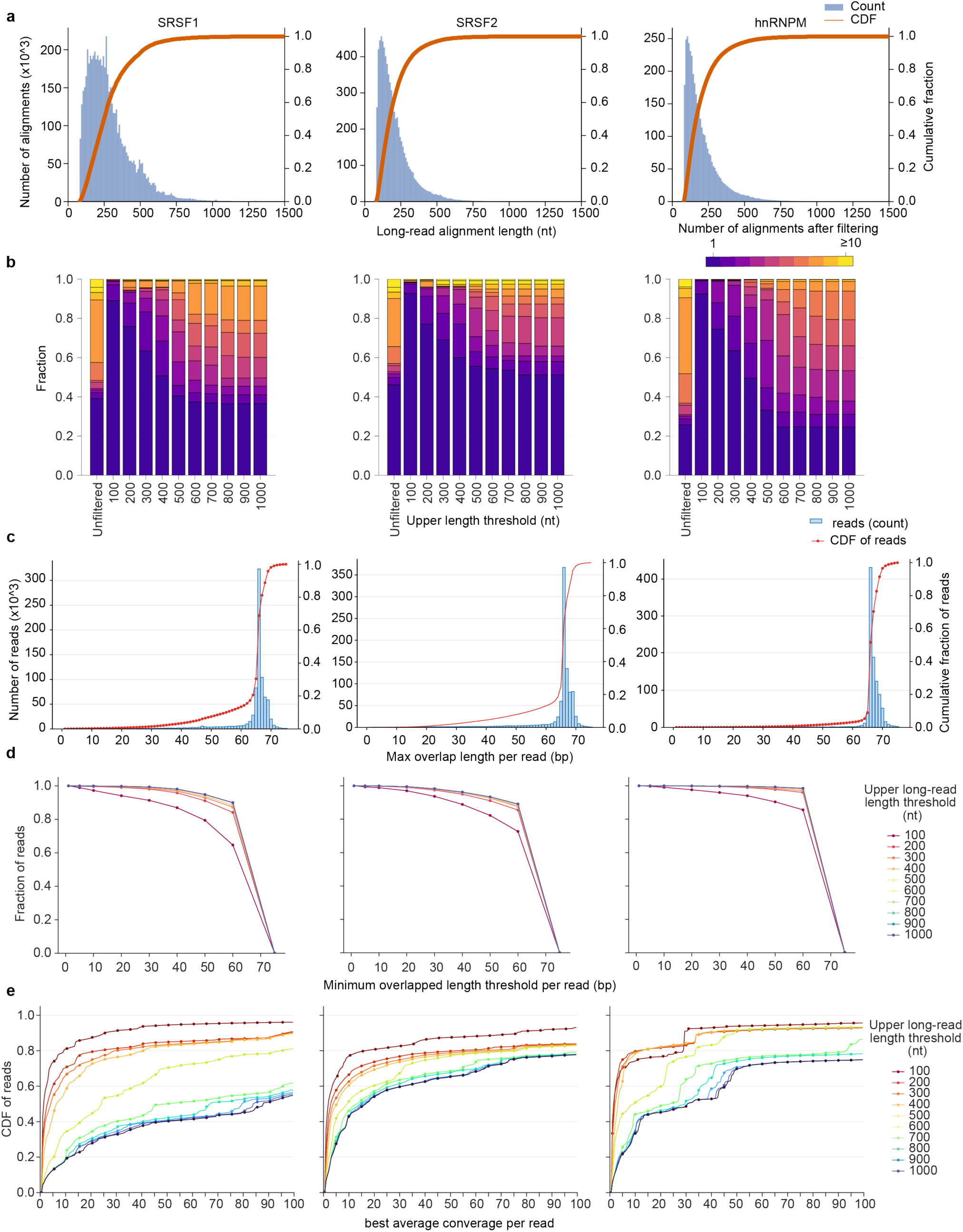
Thresholding strategy for long-read-guided alignment selection a–e, Analyses were performed for SRSF1, SRSF2 and hnRNPM for LRA parameter calibration. **a,** Distributions and cumulative fractions of aligned lengths for long-read CLIP. Aligned length was defined as the sum of BED12 block sizes (intronic segments excluded). **b,** Stacked bar plots showing the distribution of the number of candidate alignments per read before and after long-read-guided filtering using mask overlap and splice-junction consistency for each upper length threshold. **c,** Distributions and cumulative fractions of the maximum mask-overlap length per short read. For each short read, mask-overlap length was computed for each candidate alignment as the number of overlapped nucleotides with the corresponding long-read-derived mask, and the maximum value across candidate alignments for that read was used. **d,** Fraction of short reads that retain at least one candidate alignment with mask-overlap length exceeding each overlap threshold (x-axis). Colors indicate the upper bound on long-read mapping length used when generating the long-read-derived mask. **e,** Cumulative fraction of the per-read maximum long-read support score (defined as the mean long-read depth across genomic positions spanned by each candidate alignment; maximum across candidate alignments per read). Colors indicate the same long-read mapping-length upper bound as in **d**.

**Extended Data Fig. 3.**
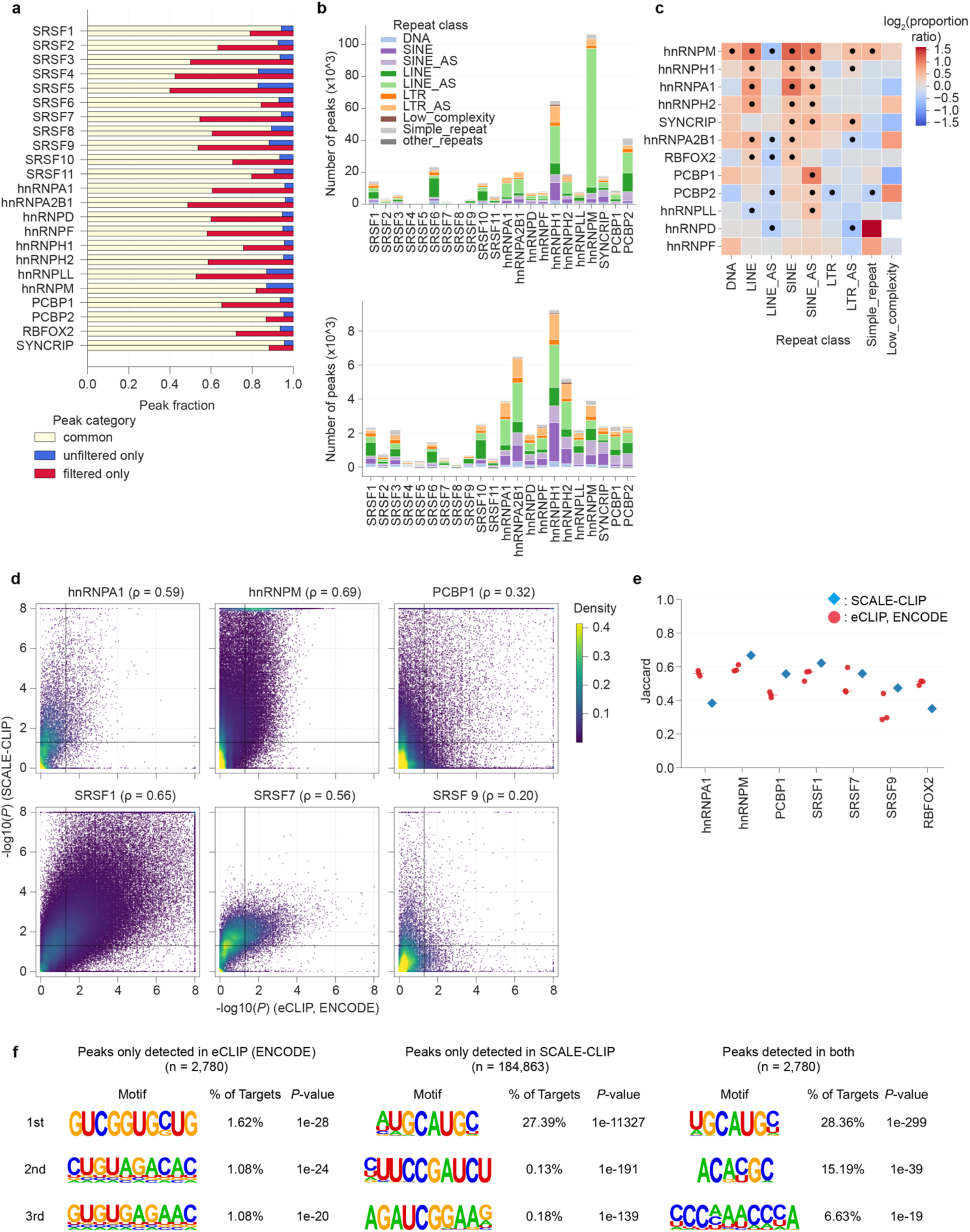
Peak characteristics and specificity of SCALE-CLIP. **a,** Fraction of peaks filtered out and newly detected after long-read-guided filtering of short-read alignments for each RBP. **b,** Composition of repeat-overlapping peaks across RepeatMasker classes for each RBP (top). Change in the number of detected repeat-overlapping peaks after long-read-guided filtering (filtered minus unfiltered) for each repeat class (bottom). **c,** Heatmap showing repeat-class composition differences between filtered-only and unfiltered-only peaks for each RBP. Color indicates the log2 ratio of the proportion of each RepeatMasker class in filtered-only versus unfiltered-only peaks within each RBP. Positive values indicate relative enrichment in filtered-only peaks, and negative values indicate relative enrichment in unfiltered-only peaks. Black dots indicate classes with significant compositional bias by two-sided Fisher’s exact test followed by Benjamini-Hochberg correction (q < 0.05). Cells with fewer than 20 total peaks across the two peak sets were masked. **d,** Density-encoded scatter plot comparing window-level CLIP enrichment between ENCODE eCLIP (x-axis) and SCALE-CLIP (y-axis) for seven benchmark SFs. Each point represents a window tested in both datasets; color intensity indicates local point density estimated by fastKDE. Enrichment is defined per window as the maximum −log₁₀ (*P*) for IP-versus-input enrichment across replicates. **e,** Reproducibility of window-level enrichment across replicates was assessed by Jaccard similarity between enriched-window sets (q-value < 0.2), computed on the intersection of tested windows between replicates. For SCALE-CLIP (three replicates), all pairwise values are shown with the median; for ENCODE (two replicates), the single replicate-pair value is shown. **f,** De novo motif analysis of RBFOX2 peaks classified as ENCODE eCLIP-only, SCALE-CLIP-only, or common to both datasets. For each peak class, the top three enriched motifs are shown with the percentage of target sequences containing the motif and the corresponding enrichment *P*-value. Numbers in parentheses indicate the number of peaks in each category.

**Extended Data Fig. 4.**
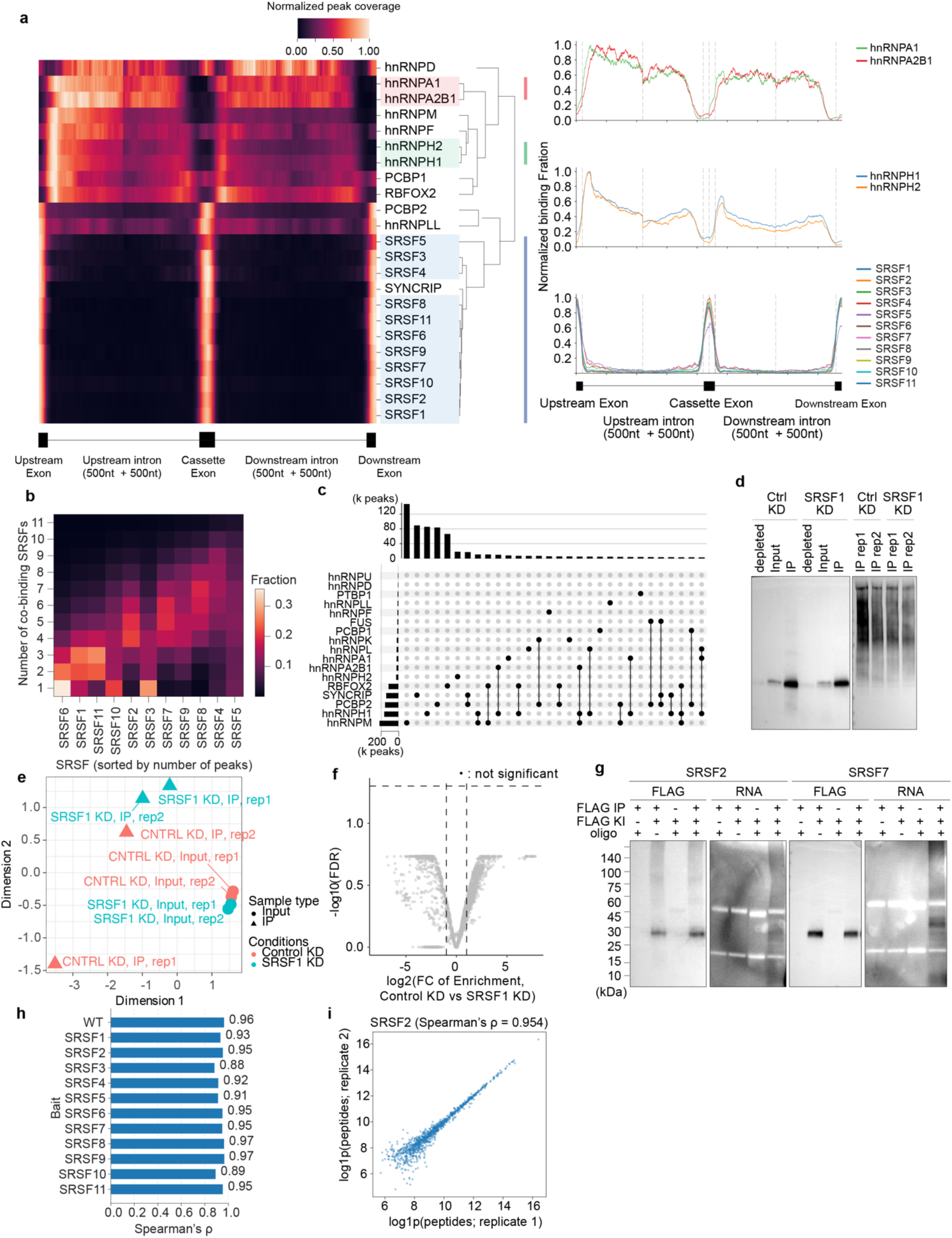
SCALE-CLIP binding architecture and co-binding characteristics. **a,** Heatmap showing meta-exon binding profiles for each profiled RBP, computed by aggregating fine-mapped SCALE-CLIP peaks around rMATS-defined cassette exons (SE) with ± 50 nt exonic and ± 500 nt intronic flanks. Values were scaled within each RBP (row-wise max = 1) to emphasize binding architecture, and RBPs were clustered by similarity of their meta-exon profiles. Line plots show the average SCALE-CLIP meta-exon profiles for representative RBP clusters (right). **b,** Fraction of peaks stratified by the number of co-binding factors for each SRSF. **c,** UpSet plots showing the shared binding peaks among hnRNP family members. Each bar represents the number of peaks shared by the indicated combinations of factors. **d,** Anti-FLAG immunoblot (left) and detection of recovered biotinylated RNA fragments (right) in control and SRSF1 knockdown samples in SRSF7-3×FLAG cells for validation of KD-CLIP-seq sample preparation. **e,** Multidimensional scaling (MDS) plot of SRSF7 CLIP-seq samples in **d**, showing separation by assay (IP vs input) and condition (control vs SRSF1 KD) as indicated. **f,** Volcano plot of differential SRSF7 IP/Input enrichment (edgeR generalized linear model). X- and y-axes indicate log2 fold change and -log10 FDR, respectively. Significance thresholds are FDR < 0.1 and |log2 fold change of enrichment| > 1. No sites passed both thresholds. **g,** RNA oligonucleotide binding assay results for SRSF2 and SRSF7. Membranes were probed with anti-FLAG antibody, and biotinylated RNA was visualized using avidin-HRP in reactions under the indicated conditions (± FLAG IP, ± RNA oligos and lysates from FLAG-tagged or parental cells). **h,** Spearman’s ρ between IP-MS replicates for each bait. **i,** Representative replicate scatter plot for SRSF2 IP-MS. Peptide counts from the two replicate immunoprecipitations are shown after log1p transformation.

**Extended Data Fig. 5.**
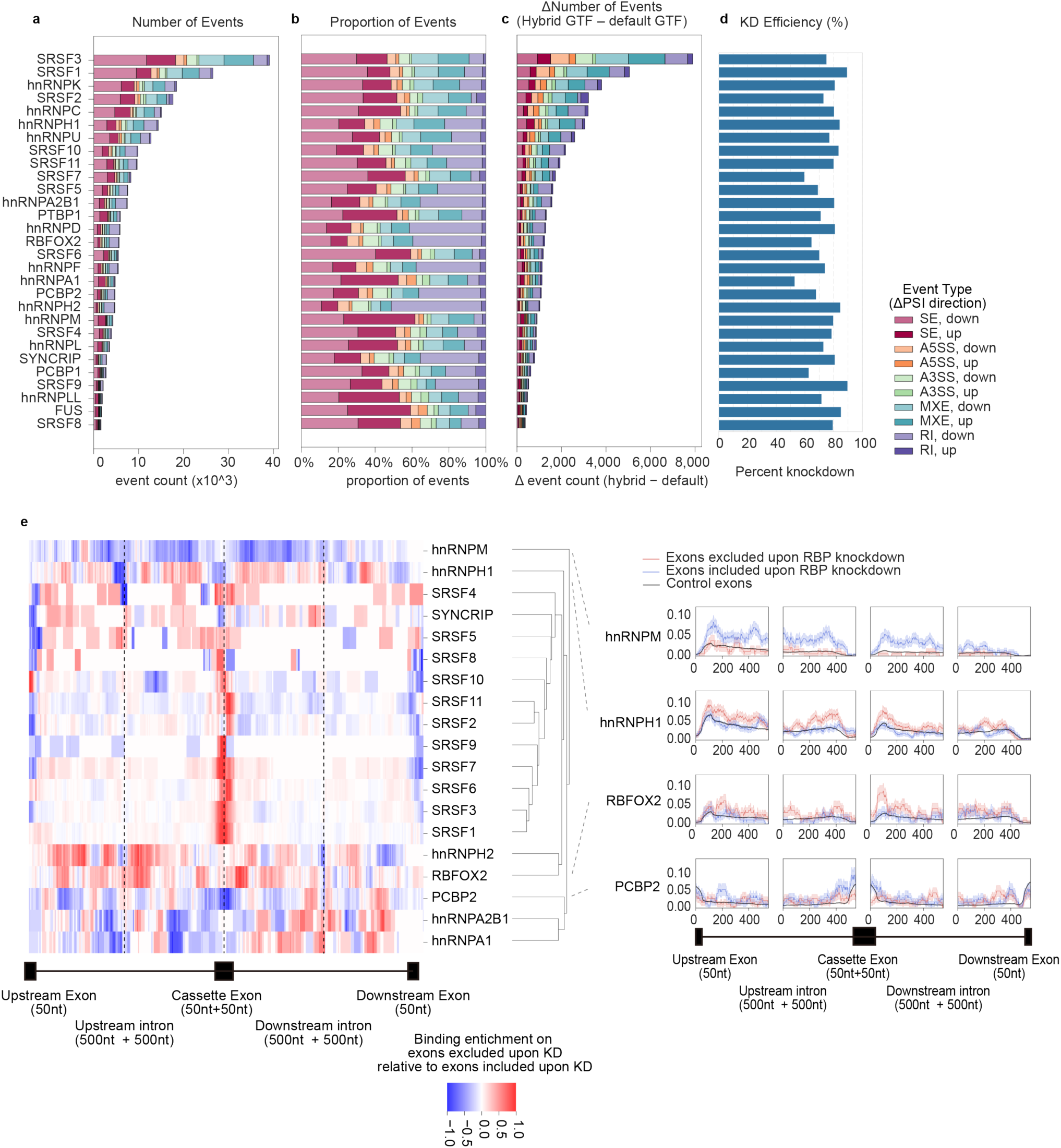
Splicing summary statistics and expanded positional binding maps supporting bidirectional positional regulatory rules of SRSFs. Each panel summarizes alternative splicing statistics for RNA-seq datasets following individual RBP knockdowns. **a,** Bar plot indicating the number of significant splicing alteration events (|ΔPSI| > 5%, FDR < 0.05) for each type of alternative splicing event. **b,** Proportions of different alternative splicing event types among significant changes. **c,** Bar plot showing the difference in the number of significant splicing alteration events (|ΔPSI| > 5% and FDR < 0.05) detected by rMATS-turbo when using the default GENCODE v44 annotation GTF versus the hybrid-assembled GTF, grouped by event type. Difference was calculated as hybrid-assembled GTF minus default GENCODE v44 GTF. **d,** Knockdown efficiency of each targeted RBP, represented as the percent reduction in TMM-normalized average read counts relative to control samples across replicates. **e,** Expanded view of the positional binding signatures shown in Fig. 3a. Left, heatmap of differential SCALE-CLIP peak-density profiles around cassette exons excluded versus included upon knockdown for each RBP, calculated as Δbinding = density (excluded) – density (included) using cassette exons with |ΔPSI| > 5% and FDR < 0.05 in matched KD RNA-seq. RBPs were hierarchically clustered by similarity of Δbinding profiles. Right, mean meta-exon SCALE-CLIP profiles for representative non-SRSF RBPs, plotted for exons excluded upon knockdown (red), included upon knockdown (blue), and control exons (black).

**Extended Data Fig. 6.**
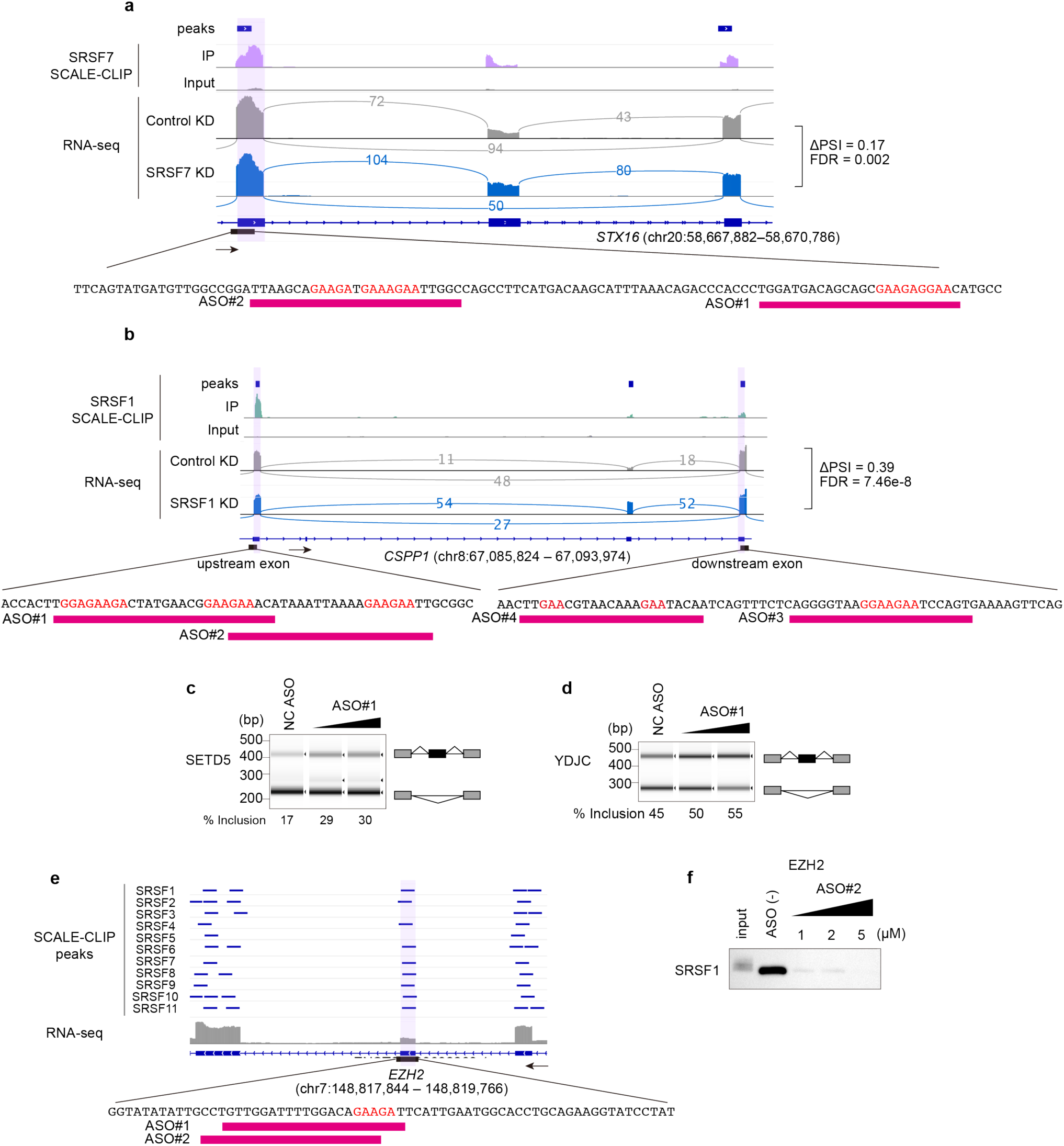
Additional ASO validations and target maps supporting the SRSF positional code a, b, Expanded views of the STX16 (a) and CSPP1 (b) loci shown in Fig. 3c,d. Upper tracks show SRSF7 SCALE-CLIP peaks (a) and SRSF1 SCALE-CLIP peaks (b).Lower tracks show short-read RNA-seq coverage in control versus SRSF7 knockdown (a) and control versus SRSF1 knockdown (b). Magenta bars indicate ASO target regions. Red letters denote GAAGA-containing SRSF motif instances. **c,** Expanded view of the EZH2 poison exon shown in Fig. 3h. Multiple SRSF family proteins bind this region, and two ASOs were designed against the indicated GAAGA-containing motifs (magenta bars). **d, e,** RT-PCR validation of SETD5 (d) and YDJC (e) splicing after transfection of an ASO targeting the SCALE-CLIP-defined SRSF binding sites, compared with a non-targeting control ASO. ASO#1 was tested at increasing doses (0.05 and 0.1µM; non-targeting control, 0.1µM) for 24h. **f,** RNA oligo pull-down followed by anti-FLAG immunoblot showing dose-dependent binding of SRSF1-3×FLAG to a biotinylated RNA oligo derived from the EZH2 alternative exon in the presence of increasing concentrations of competitor ASO#2 (1, 2, or 5 μM), consistent with ASO-mediated blockade of the SRSF-binding site.

**Extended Data Fig. 7.**
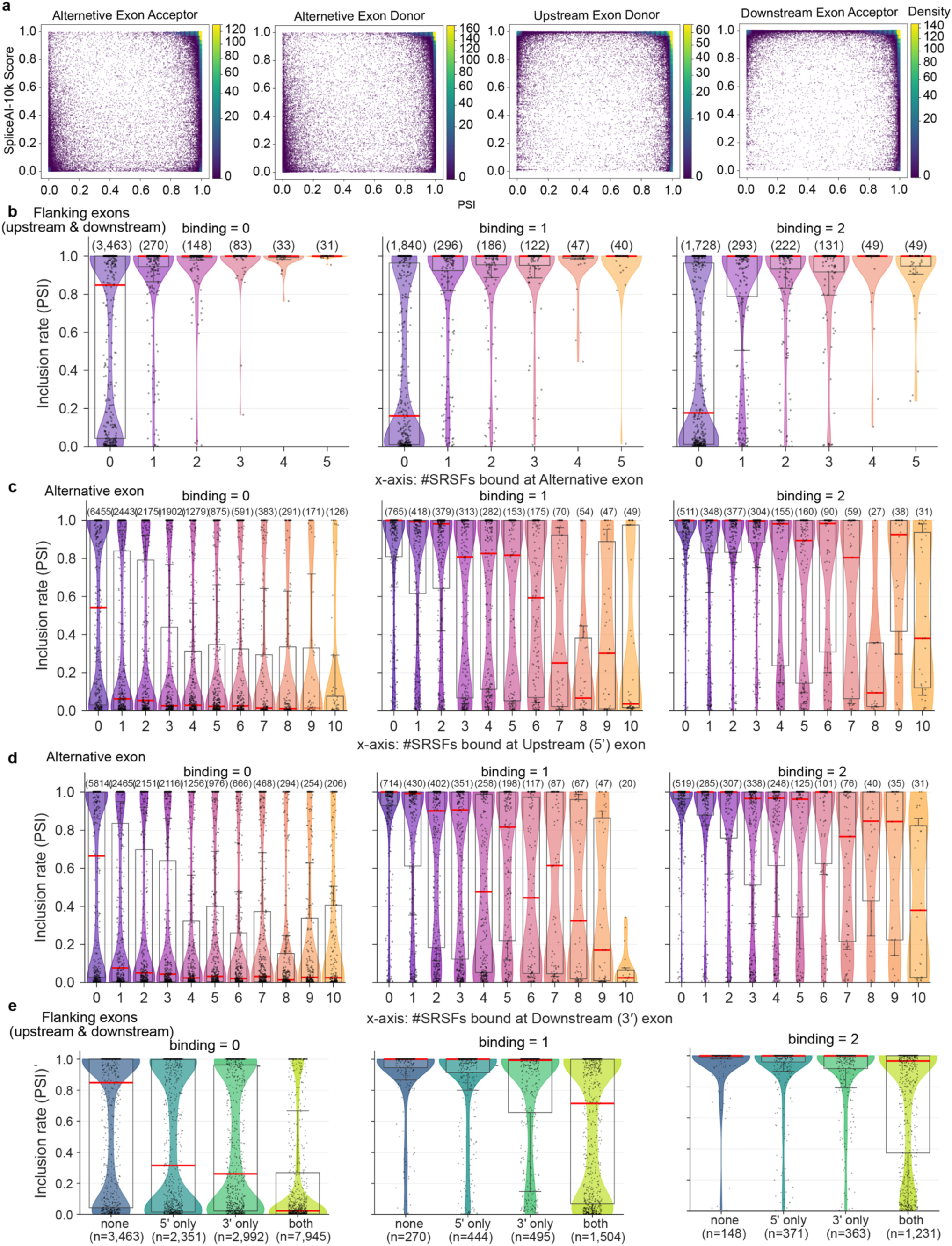
Exon inclusion as a function of splice-site strength and SRSF binding multiplicity on alternative and flanking exons. **a,** Density-encoded scatter showing the relationship between SpliceAI-10k splice site scores and exon inclusion. Each point represents an alternative exon. The x-axis indicates SpliceAI scores for donor/acceptor sites of alternative exons and their flanking upstream and downstream exons, and the y-axis indicates mean PSI across control replicates. Local data density is visualized by fastKDE with a power-law normalization (γ = 0.5) and percentile clipping (p1-p40) to enhance visualization. **b,** PSI distribution for alternative exons with suboptimal splice sites (SpliceAI-10k < 0.6 for both donor and acceptor sites), stratified by SRSF binding multiplicity on the alternative exon (x-axis), further stratified by SRSF binding multiplicity on the flanking exons. Points represent individual exons (jittered), overlaid on boxplots (center line, median; box, interquartile range; whiskers, 1.5 × IQR). Alternative exons were subselected based on the total SRSF binding number across the upstream and downstream flanking exons, and PSI distributions are shown for representative multiplicity bins (0, 3, 6, or 10 overlaps). **c, d,** PSI distributions for alternative exons with suboptimal splice sites (defined as max(SpliceAI-10k donor, acceptor) < 0.7), plotted as a function of SRSF binding multiplicity on the upstream flanking exon (c) or downstream flanking exon (d) (x-axis). Exons are further stratified by SRSF binding multiplicity on the alternative exon body (0, 1, or 2; shown as separate columns). Each point represents an individual exon (jittered). Boxplots show the median (center line), interquartile range (box), and whiskers extending to 1.5 × IQR. **e,** PSI distributions of alternative exons with suboptimal splice sites (max SpliceAI-10k score < 0.7) grouped by SRSF binding scores on upstream, downstream and both exons, stratified by binding scores on flanking exons (0, 1, 2 or 3). Group sizes are shown as the number of alternative exons in c–e and of significant splicing events in b (displayed as *n* above or under boxplots).

**Extended Data Fig. 8.**
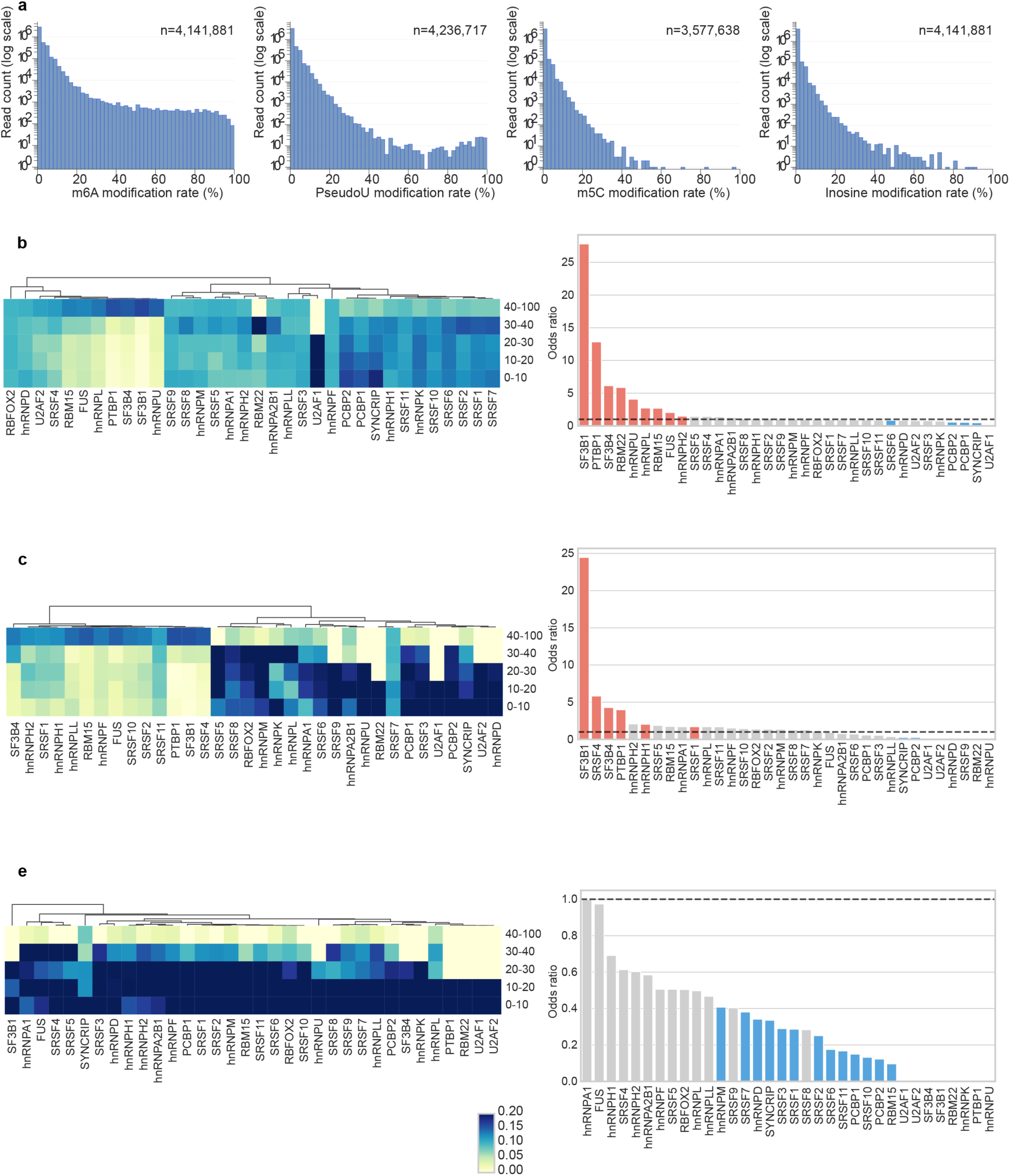
Splicing factor binding enrichment at RNA modified bases. Distribution and enrichment of SF binding relative to m^5^C, pseudouridine, and inosine RNA modifications. **a,** Per-base modification frequency distributions for m^6^A, pseudouridine, m^5^C and inosine detected using Dorado. Each histogram shows the number of bases (y-axis) within each modification frequency bin (x-axis), defined as (Nmod / Nvalid_cov) x 100. Nmod and Nvalid_cov represent the number of reads called as modified and the total number of valid reads considered for modification-frequency estimation for each modification, respectively. **b–e,** Left is heatmap showing normalized overlap rate between RBP binding peaks and per-base modification frequency bins for m^6^A (b), pseudouridine (c), m^5^C (d) and inosine (e). RBPs were hierarchically clustered using Euclidean distance and complete linkage. Modification bins for pseudouridine, m^5^C and inosine were limited to five ranges (0-10%, 10-20%, 20-30%, 30-40% and ≥ 40%) due to sparse high-frequency bases. Right shows enrichment of RBP binding at highly modified bases. Bars show odds ratios (OR) for peak overlap of high-modification sites (≥ 40%) versus low-modification sites (≤ 10%). Red/blue indicate significant enrichment/depletion (two-sided Fisher’s exact test, Benjamini-Hochberg FDR < 0.05); grey, not significant.

**Extended Data Fig. 9.**
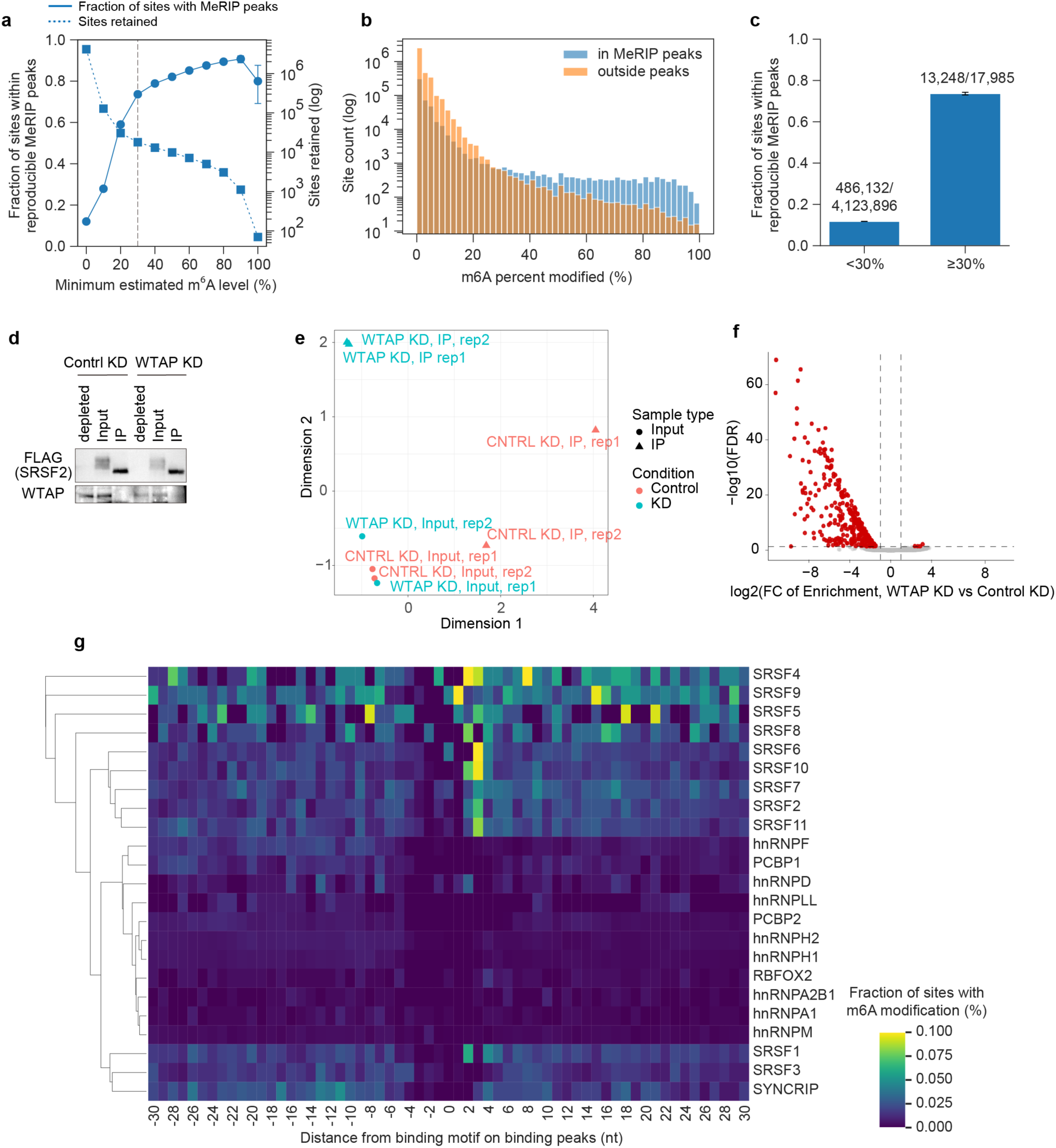
Validation and motif-centered positioning of high-confidence m^6^A sites in splicing factor binding peaks. **a,** Fraction of Dorado-called m^6^A sites that fall within reproducible MeRIP-seq peaks across increasing minimum estimated m^6^A thresholds. The number of retained sites at each threshold is shown on the right y-axis. Based on the increase in orthogonal MeRIP support while retaining a substantial number of sites, a threshold of 30% estimated modification was used to define a higher-confidence m^6^A set for downstream analyses. **b,** Histogram of estimated m^6^A levels for Dorado-called sites within versus outside reproducible MeRIP-seq peaks. Sites within reproducible MeRIP-seq peaks are shifted toward higher estimated modification levels. The vertical dashed line indicates the 30% threshold used in downstream analyses. **c,** Summary of overlap with reproducible MeRIP-seq peaks for sites with estimated m^6^A levels below 30% versus at least 30%. Error bars indicate 95% Wilson confidence intervals. **d,** Anti-FLAG and anti-WTAP immunoblot confirming WTAP knockdown and FLAG expression in control and WTAP knockdown cells. **e,** Multidimensional scaling (MDS) plot of seCLIP samples in (d), showing separation by assay (IP vs input) and condition (control vs WTAP KD) as indicated. **f,** Volcano plot showing differential binding enrichment of SRSF2 in WTAP knockdown versus control. Differential binding was estimated with DESeq2 using a GLM with an interaction term (design: ∼ assay + condition + assay:condition; assay = IP vs Input, condition = KD vs control). Axes indicate log2 fold change and -log10 FDR. Significantly altered sites were defined as BH-FDR < 0.05 and |log2FC| > 1. **g,** Motif-centered metaprofile of high-confidence m^6^A sites within RBP binding peaks. For each RBP, the top enriched sequence motif identified by HOMER was used as the anchor, with the motif center defined as position 0. The plot shows, for each offset from the motif center (−30 to +30 nt), the fraction of motif hits that contain a high-confidence m^6^A site at that offset. High-confidence m^6^A sites were defined as sites with modification frequency ≥ 30% in Nanopore direct RNA sequencing, read depth ≥ 20, and an A-call fraction ≥ 0.7. At each offset, the fraction was computed as (number of motif hits with an m^6^A at that offset) / (total number of motif hits). Offsets were computed in a strand-aware manner based on motif orientation, and only m^6^A sites overlapping the peak on the same strand were considered. For even-length motifs, position 0 was defined as the downstream of the two central nucleotides.

**Extended Data Fig. 10.**
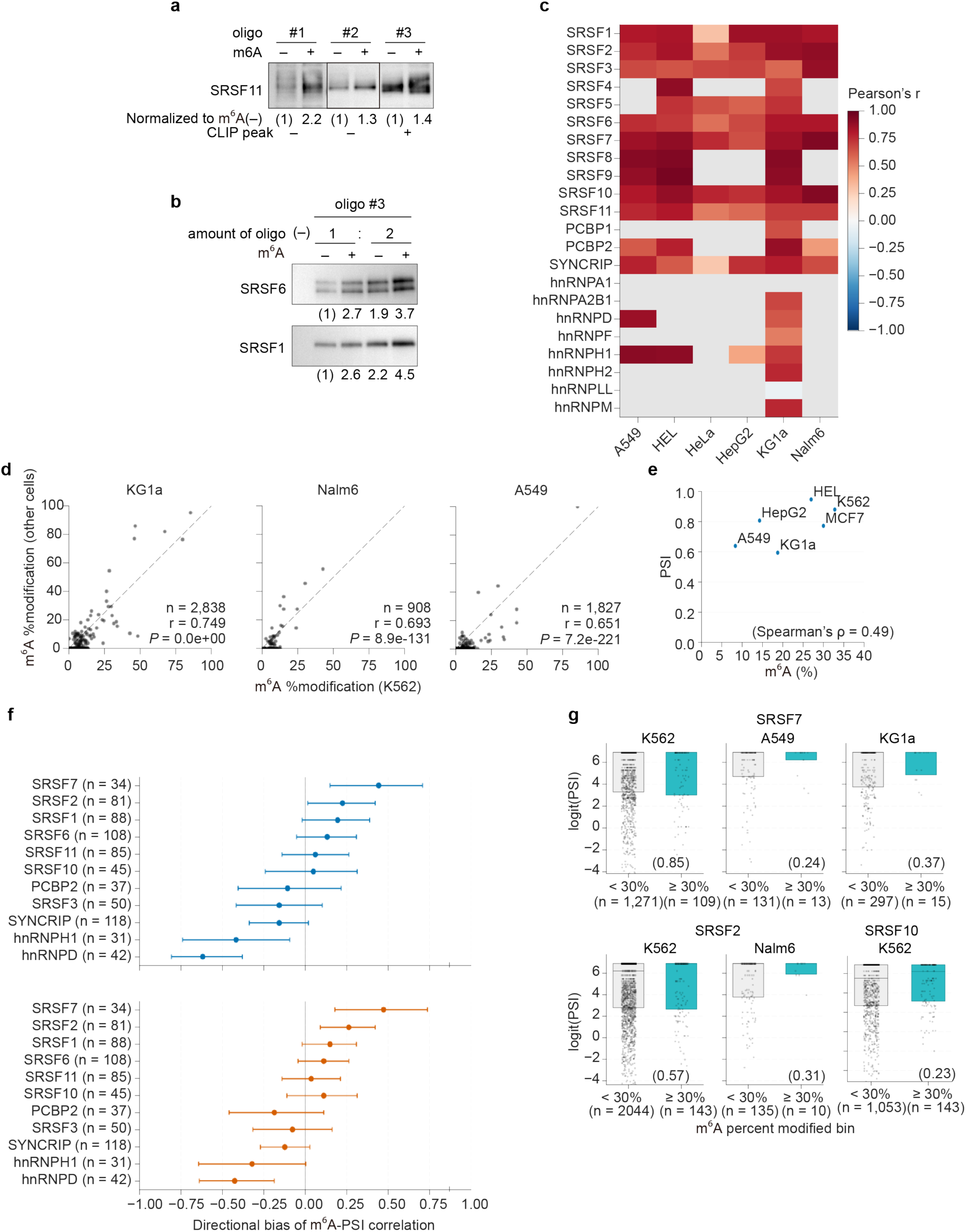
m^6^A-dependent enhancement of SRSF binding and cross-cell-type concordance of m^6^A-associated splicing regulation. **a,** Biotinylated RNA oligos (20–30 nt) were synthesized as matched A versus m^6^A pairs (Supplementary Data 6) and incubated with lysates expressing 3×FLAG-tagged SRSF11, followed by streptavidin pull-down and anti-FLAG immunoblot. A representative experiment is shown. Panels were imaged with exposure optimized per panel; band intensities should be compared within each matched A/m^6^A pair. Quantification of binding to the m^6^A-containing oligo normalized to the corresponding unmodified A oligo is shown below each blot, together with an annotation indicating whether the corresponding locus is detected as a CLIP peak in SCALE-CLIP. **b,** Titration of input oligo amount for oligo #3. Pull-down assays were performed for SRSF1 and SRSF6 using two input oligo amounts (1× and 2×). Immunoblots show the recovered SRSF signal, and the binding to the m^6^A-containing oligo normalized to the unmodified (A) oligo is shown below. **c,** Heatmap showing cross-cell-type concordance of m^6^A levels at sites overlapping K562-defined RBP peaks. For each RBP, Pearson correlation coefficients were calculated between percent m^6^A in K562 and in the indicated cell type across overlapping sites. Gray cells indicate ≤ 20 overlapping sites, for which values are not shown. **d,** Representative example for (c) showing percent m^6^A values at sites overlapping SRSF1 peaks, comparing K562 (x-axis) with KG1a, Nalm6, and A549 (y-axis). **e,** Example locus-level concordance of the m^6^A-PSI relationship across cell types. Scatter plot shows, across cell types, the association between percent m^6^A at a specific SRSF1-overlapping site and cassette-exon inclusion (PSI) measured in the matched RNA-seq dataset. **f,** Bias in the direction of m^6^A-PSI association across cell types, stratified by overlap with each RBP’s binding peaks. Distributions show Spearman’s ρ values for eligible m^6^A sites. Only sites with sufficient m^6^A coverage and matched PSI estimates in ≥ 4 cell types were included. The top panel includes all eligible sites, whereas the bottom panel restricts to strongly associated sites (|ρ| > 0.2). **g,** Cell-type-specific PSI distributions for cassette exons linked to SRSF2, 7 and 10-overlapping m^6^A sites, stratified by m^6^A level at the corresponding site (<30% or ≥30%). Points represent individual pairs; boxplots summarize the proposed relationship among m^6^A, effective SRSF occupancy and splicing outcomes. Boxplots show the median, interquartile range and 1.5 × IQR. Group sizes indicate the number of cassette exon-m^6^A site pairs. *P* values from two-sided Mann–Whitney U tests are shown in parentheses.

## Supplementary Data

Supplementary Data 1. Peak-enrichment comparison between ENCODE eCLIP and SCALE-CLIP datasets.

Contains RBP-specific peak-level enrichment statistics used to compare ENCODE eCLIP and SCALE-CLIP datasets, including minimum enrichment P values and corresponding −log10(*P*) values used in Fig. 1d and Extended Data Fig. 3d.

Supplementary Data 2. Master table of alternative splicing events. Contains rMATS-turbo alternative splicing events annotated with exon coordinates, splice-site scores, SRSF CLIP peak overlaps and splicing responses after individual SRSF knockdown. Transcript-oriented 5′ and 3′ exon columns indicate flanking exons relative to the alternative exon.

Supplementary Data 3. RNA modification–RBP binding overlap by modification-level bins. Contains binned overlap summaries between RNA modification sites and RBP CLIP peaks, including the number of sites, number of overlapping sites, overlap fraction and heatmap values for each modification type and RBP, used for Fig. 5a and Extended Data Fig. 8b–d.

Supplementary Data 4. IP-MS quantification matrix. Contains protein-level IP-MS quantification values for parental control and tagged cell lines. Rows represent quantified protein groups with UniProt accessions, gene symbols and UniProt entry names, and columns report replicate-level protein intensities.

Supplementary Data 5. Differential CLIP binding statistics after knockdown. Contains differential CLIP binding statistics for CLIP peaks after knockdown. Rows report genomic coordinates, peak identifiers, strand, GC content, log2 fold change, average log counts per million, F statistic, nominal P value and FDR-adjusted significance.

Supplementary Data 6. Sequences of guides, donors, siRNAs, primers, RNA oligos and ASOs. Contains sequences of reagents used in this study, including sgRNAs and ssODN donors for knock-in cell-line generation, siRNAs and DsiRNAs for knockdown, PCR/qPCR/RT-PCR primers, RNA oligonucleotides and antisense oligonucleotides.

Supplementary Data 7. External datasets and resources. Contains accession identifiers and metadata for public datasets and external resources used in comparative analyses, including ENCODE eCLIP files and additional public datasets.

## References

1. Nilsen, T. W. & Graveley, B. R. Expansion of the eukaryotic proteome by alternative splicing. Nature 463, 457–463 (2010).

2. Dvinge, H., Kim, E., Abdel-Wahab, O. & Bradley, R. K. RNA splicing factors as oncoproteins and tumour suppressors. Nat Rev Cancer 16, 413–430 (2016).

3. Scotti, M. M. & Swanson, M. S. RNA mis-splicing in disease. Nat Rev Genet 17, 19–32 (2016).

4. Shine, M. et al. Co-transcriptional gene regulation in eukaryotes and prokaryotes. Nat Rev Mol Cell Biol 1–21 (2024) doi:10.1038/s41580-024-00706-2.

5. Ule, J. & Blencowe, B. J. Alternative Splicing Regulatory Networks: Functions, Mechanisms, and Evolution. Molecular Cell 76, 329–345 (2019).

6. Wang, Z. & Burge, C. B. Splicing regulation: From a parts list of regulatory elements to an integrated splicing code. RNA 14, 802–813 (2008).

7. Martinez, N. M. et al. Pseudouridine synthases modify human pre-mRNA co-transcriptionally and affect pre-mRNA processing. Molecular Cell 82, 645–659.e9 (2022).

8. Mendel, M. et al. Splice site m6A methylation prevents binding of U2AF35 to inhibit RNA splicing. Cell 184, 3125–3142.e25 (2021).

9. Xiao, W. et al. Nuclear m 6 A Reader YTHDC1 Regulates mRNA Splicing. Molecular Cell 61, 507–519 (2016).

10. Hafner, M. et al. Transcriptome-wide Identification of RNA-Binding Protein and MicroRNA Target Sites by PAR-CLIP. Cell 141, 129–141 (2010).

11. Hafner, M. et al. CLIP and complementary methods. Nat Rev Methods Primers 1, 1–23 (2021).

12. König, J. et al. iCLIP reveals the function of hnRNP particles in splicing at individual nucleotide resolution. Nat Struct Mol Biol 17, 909–915 (2010).

13. Licatalosi, D. D. et al. HITS-CLIP yields genome-wide insights into brain alternative RNA processing. Nature 456, 464–469 (2008).

14. Van Nostrand, E. L. et al. Robust transcriptome-wide discovery of RNA-binding protein binding sites with enhanced CLIP (eCLIP). Nat Methods 13, 508–514 (2016).

15. Xiang, J. S., Danielle M. Schafer, Rothamel, K. L. & Yeo, G. W. Decoding protein–RNA interactions using CLIP-based methodologies. Nat Rev Genet 25, 879–895 (2024).

16. Karni, R. et al. The gene encoding the splicing factor SF2/ASF is a proto-oncogene. Nat Struct Mol Biol 14, 185–193 (2007).

17. Anczuków, O. et al. SRSF1-Regulated Alternative Splicing in Breast Cancer. Molecular Cell 60, 105–117 (2015).

18. Leclair, N. K. et al. Poison Exon Splicing Regulates a Coordinated Network of SR Protein Expression during Differentiation and Tumorigenesis. Molecular Cell 80, 648–665.e9 (2020).

19. Van Nostrand, E. L. et al. A large-scale binding and functional map of human RNA-binding proteins. Nature 583, 711–719 (2020).

20. Feng, H. et al. Modeling RNA-Binding Protein Specificity In Vivo by Precisely Registering Protein-RNA Crosslink Sites. Molecular Cell 74, 1189–1204.e6 (2019).

21. Treangen, T. J. & Salzberg, S. L. Repetitive DNA and next-generation sequencing: computational challenges and solutions. Nat Rev Genet 13, 36–46 (2012).

22. Zhang, Z. & Xing, Y. CLIP-seq analysis of multi-mapped reads discovers novel functional RNA regulatory sites in the human transcriptome. Nucleic Acids Research 45, 9260–9271 (2017).

23. Paul, S. et al. RNA molecules display distinctive organization at nuclear speckles. iScience 27, 109603 (2024).

24. Spector, D. L. & Lamond, A. I. Nuclear Speckles. Cold Spring Harbor Perspectives in Biology 3, a000646–a000646 (2011).

25. Boyle, E. A. et al. Skipper analysis of eCLIP datasets enables sensitive detection of constrained translation factor binding sites. Cell Genomics 3, 100317 (2023).

26. Zhou, D., Couture, S., Scott, M. S. & Abou Elela, S. RBFOX2 alters splicing outcome in distinct binding modes with multiple protein partners. Nucleic Acids Research 49, 8370–8383 (2021).

27. Busch, A. & Hertel, K. J. Evolution of SR protein and hnRNP splicing regulatory factors. WIREs RNA 3, 1–12 (2012).

28. Yee, B. A., Pratt, G. A., Graveley, B. R., Van Nostrand, E. L. & Yeo, G. W. RBP-Maps enables robust generation of splicing regulatory maps. RNA 25, 193–204 (2019).

29. Grammatikakis, I. et al. Alternative Splicing of Neuronal Differentiation Factor TRF2 Regulated by HNRNPH1/H2. Cell Reports 15, 926–934 (2016).

30. Harvey, S. E. et al. Coregulation of alternative splicing by hnRNPM and ESRP1 during EMT. RNA 24, 1326–1338 (2018).

31. Huelga, S. C. et al. Integrative Genome-wide Analysis Reveals Cooperative Regulation of Alternative Splicing by hnRNP Proteins. Cell Reports 1, 167–178 (2012).

32. Weyn-Vanhentenryck, S. M. et al. HITS-CLIP and Integrative Modeling Define the Rbfox Splicing-Regulatory Network Linked to Brain Development and Autism. Cell Reports 6, 1139–1152 (2014).

33. Shen, H., Kan, J. L. C. & Green, M. R. Arginine-Serine-Rich Domains Bound at Splicing Enhancers Contact the Branchpoint to Promote Prespliceosome Assembly. Molecular Cell 13, 367–376 (2004).

34. Yoshimi, A. et al. Coordinated alterations in RNA splicing and epigenetic regulation drive leukaemogenesis. Nature 574, 273–277 (2019).

35. Lewis, C. J. T., Pan, T. & Kalsotra, A. RNA modifications and structures cooperate to guide RNA–protein interactions. Nat Rev Mol Cell Biol 18, 202–210 (2017).

36. Deschamps-Francoeur, G., Simoneau, J. & Scott, M. S. Handling multi-mapped reads in RNA-seq. Computational and Structural Biotechnology Journal 18, 1569–1576 (2020).

37. Zhang, Z. & Xing, Y. CLIP-seq analysis of multi-mapped reads discovers novel functional RNA regulatory sites in the human transcriptome. Nucleic Acids Research 45, 9260–9271 (2017).

38. Änkö, M.-L. et al. The RNA-binding landscapes of two SR proteins reveal unique functions and binding to diverse RNA classes. Genome Biol 13, R17 (2012).

39. Pandit, S. et al. Genome-wide Analysis Reveals SR Protein Cooperation and Competition in Regulated Splicing. Molecular Cell 50, 223–235 (2013).

40. Witten, J. T. & Ule, J. Understanding splicing regulation through RNA splicing maps. Trends in Genetics 27, 89–97 (2011).

41. Llorian, M. et al. Position-dependent alternative splicing activity revealed by global profiling of alternative splicing events regulated by PTB. Nat Struct Mol Biol 17, 1114–1123 (2010).

42. Lovci, M. T. et al. Rbfox proteins regulate alternative mRNA splicing through evolutionarily conserved RNA bridges. Nat Struct Mol Biol 20, 1434–1442 (2013).

43. Ule, J. et al. An RNA map predicting Nova-dependent splicing regulation. Nature 444, 580–586 (2006).

44. Hua, Y., Vickers, T. A., Okunola, H. L., Bennett, C. F. & Krainer, A. R. Antisense Masking of an hnRNP A1/A2 Intronic Splicing Silencer Corrects SMN2 Splicing in Transgenic Mice. The American Journal of Human Genetics 82, 834–848 (2008).

45. Sinha, R. et al. Antisense oligonucleotides correct the familial dysautonomia splicing defect in IKBKAP transgenic mice. Nucleic Acids Research 46, 4833–4844 (2018).

46. Chen, C., Zhao, X., Kierzek, R. & Yu, Y.-T. A Flexible RNA Backbone within the Polypyrimidine Tract Is Required for U2AF^65^ Binding and Pre-mRNA Splicing *In Vivo*. Molecular and Cellular Biology 30, 4108–4119 (2010).

47. An, S. et al. Integrative network analysis identifies cell-specific trans regulators of m6A. Nucleic Acids Research 48, 1715–1729 (2020).

48. Liu, N. et al. N6-methyladenosine-dependent RNA structural switches regulate RNA–protein interactions. Nature 518, 560–564 (2015).

49. Blue, S. M. et al. Transcriptome-wide identification of RNA-binding protein binding sites using seCLIP-seq. Nat Protoc 17, 1223–1265 (2022).

50. Jiang, H., Lei, R., Ding, S.-W. & Zhu, S. Skewer: a fast and accurate adapter trimmer for next-generation sequencing paired-end reads. BMC Bioinformatics 15, 182 (2014).

51. Chen, S., Zhou, Y., Chen, Y. & Gu, J. fastp: an ultra-fast all-in-one FASTQ preprocessor. Bioinformatics 34, i884–i890 (2018).

52. Dobin, A. et al. STAR: ultrafast universal RNA-seq aligner. Bioinformatics 29, 15–21 (2013).

53. Li, H. Minimap2: pairwise alignment for nucleotide sequences. Bioinformatics 34, 3094–3100 (2018).

54. Choi, Y. et al. Time-resolved profiling of RNA binding proteins throughout the mRNA life cycle. Molecular Cell 84, 1764–1782.e10 (2024).

55. Chen, Y., Chen, L., Lun, A. T. L., Baldoni, P. L. & Smyth, G. K. edgeR v4: powerful differential analysis of sequencing data with expanded functionality and improved support for small counts and larger datasets. Nucleic Acids Research 53, gkaf018 (2025).

56. Dou, X. et al. RBFOX2 recognizes N6-methyladenosine to suppress transcription and block myeloid leukaemia differentiation. Nat Cell Biol 25, 1359–1368 (2023).

57. Shumate, A., Wong, B., Pertea, G. & Pertea, M. Improved transcriptome assembly using a hybrid of long and short reads with StringTie. PLoS Comput Biol 18, e1009730 (2022).

58. Pardo-Palacios, F. J. et al. SQANTI3: curation of long-read transcriptomes for accurate identification of known and novel isoforms. Nat Methods 21, 793–797 (2024).

59. Wang, Y. et al. rMATS-turbo: an efficient and flexible computational tool for alternative splicing analysis of large-scale RNA-seq data. Nat Protoc 10.1038/s41596-023-00944-2 (2024) doi:10.1038/s41596-023-00944-2.

60. Rogalska, M. E. et al. Transcriptome-wide splicing network reveals specialized regulatory functions of the core spliceosome. Science 386, 551–560 (2024).

61. Jaganathan, K. et al. Predicting Splicing from Primary Sequence with Deep Learning. Cell 176, 535–548.e24 (2019).

